# The tumour suppressor LACTB remodels mitochondria to promote cytochrome c release and apoptosis

**DOI:** 10.1101/2025.04.12.648505

**Authors:** Sukrut C. Kamerkar, Taewook Kang, Radu V. Stan, Edward J. Usherwood, Henry N. Higgs

## Abstract

Mitochondria are pivotal regulators of cellular homeostasis, integrating energy metabolism, biosynthesis, and programmed cell death (apoptosis). During apoptosis, mitochondrial outer membrane permeabilization by BAX/BAK pores facilitates release of apoptotic factors, while the role of inner mitochondrial membrane (IMM) remodelling remains less understood. Here, we identify LACTB, a filament-forming serine protease and tumour suppressor, as a regulator of IMM dynamics during apoptosis. LACTB is required for apoptosis-induced IMM remodelling, which in turn causes increased release of cytochrome c and other mitochondrial apoptotic factors. LACTB-induced membrane remodelling is independent of OPA1 processing. Rather, purified LACTB binds and remodels cardiolipin-enriched membrane nanotubes preferentially over planar lipid membranes, suggesting a direct effect in apoptotic membrane remodelling. Intriguingly, LACTB is not required for mitochondrial shape changes induced by mitochondrial depolarization, suggesting that LACTB action is apoptosis-specific. Collectively, our findings establish LACTB as a mediator of apoptosis-induced IMM remodelling, suggesting a mechanism for tumour suppression in cancer.

## Introduction

Mitochondria are indispensable regulators of cellular life and death(*1*). Beyond their well-established role in ATP production via oxidative phosphorylation, mitochondria are hubs for the biosynthesis and metabolism of signalling molecules, lipids, and metabolites critical to maintain cellular homeostasis as well as inflammation and immune responses(*2*, *3*). Paradoxically, mitochondria are equally central to programmed cell death (apoptosis), where mitochondrial outer membrane permeabilization (MOMP) mediated by BCL-2 family proteins (notably BAX and BAK) leads to the release of factors that activate the apoptotic cascade, including caspase-activating proteins (cytochrome c), proteins inhibiting caspase inhibitors (SMAC/DIABLO and HTRA2/Omi), and proteins participating in DNA cleavage (AIF and EndoG)(*4–7*).

While significant progress has been made in understanding the mechanisms driving MOMP, less is known about the molecular players orchestrating inner mitochondrial membrane (IMM) remodelling during apoptosis(*8*). However, apoptotic IMM remodelling might be important for the release of cytochrome c, 85% of which is sequestered in cristae where it is electrostatically bound to the IMM(*8*),(*9*). Several mechanisms for apoptotic IMM remodelling have been proposed, including the actions of pro-apoptotic BH3-only proteins(*9*, *10*), effects of BAX/BAK beyond MOMP, and regulation of proteolytic processing of the IMM dynamin OPA1(*8*, *11*, *12*).

LACTB is a filament-forming serine protease localized to the mitochondrial intermembrane space (IMS)(*13*). Although its physiological role in non-cancerous cells remains unclear, LACTB is well established as a tumour suppressor in several cancer types(*14–22*). However, the mechanism underlying this tumour-suppressive activity is still under debate. One study suggests that LACTB mediates the proteolysis of mitochondrially-localized phosphatidylserine decarboxylase (PISD), thereby altering mitochondrial phospholipid synthesis(*14*). Other reports have implicated cytoplasmic functions for LACTB in tumour suppression through interactions with p53, PP1A, or components of the PI3K pathway(*15*, *16*, *20*). Additionally, LACTB overexpression has been shown to increase apoptosis(*17*), although the mechanism remains elusive.

Here, we demonstrate a direct role for LACTB in promoting the release of mitochondrial pro-apoptotic proteins via remodelling of the inner mitochondrial membrane, thereby establishing LACTB as a novel apoptotic factor.

## Results

### LACTB plays a role in apoptosis

One potential mechanism for LACTB-mediated tumour suppression is through apoptosis(*17*). To test this possibility, we induced apoptosis using staurosporine(*23*) in control or LACTB knockdown (KD) HeLa cells (Fig. 1A), and measured cell viability by the sulforhodamine B assay(*24*). LACTB KD cells display increased cell viability compared to controls at 4, 6 and 7-hrs of staurosporine treatment (Fig. 1B and C). LACTB KD causes a similar apoptotic decrease in B16-F10 cells (Fig. S1A,B).

**Figure 1.**
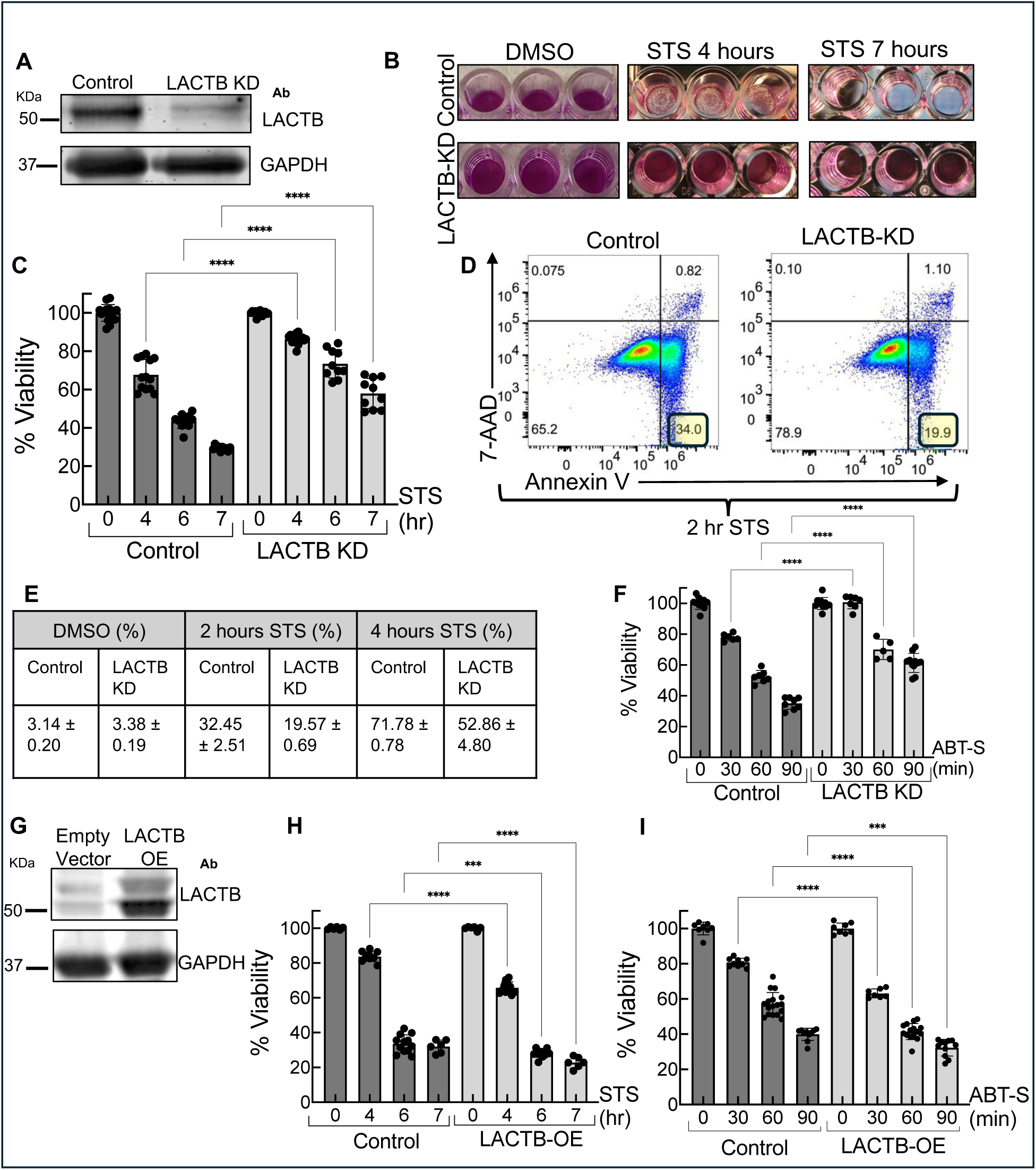
LACTB plays a role in apoptosis. (**A**) Western blot analysis of LACTB in LACTB knockdown (KD) and control HeLa cells. GAPDH serves as a loading control. (**B**) Sulforhodamine B (SRB) assay measuring cell viability in control and LACTB KD HeLa cells treated with DMSO or 1 μM staurosporine (STS) for 4 and 7 hours. (**C**) Quantification of SRB assay in control and LACTB KD cells following STS treatment (N = 3). (**D**) Flow cytometry analysis of Annexin-V and 7-AAD staining in control and LACTB KD cells treated with 1 μM STS for 2 hours. (**E**) Table quantifying the percentage of Annexin-V⁺/7-AAD⁻ cells after treatment with STS (2 and 4 hours) or DMSO (4 hours) (N = 3). (**F**) SRB assay of control and LACTB KD cells upon treatment with ABT-737 (10 μM) and S63845 (2 μM) (ABT-S) (N = 2). (**G**) Western blot analysis of LACTB in control and stable LACTB overexpressing (OE) HeLa cells. (**H**) SRB assay of control and LACTB OE cells following 1 μM STS treatment (N = 2). (I) SRB assay of control and LACTB OE cells following ABT-S treatment (N = 2). ****p < 0.0001, ***p < 0.001 (one-way ANOVA). Data are presented as mean ± SD. N indicates the number of independent experiments.

As an independent method to assess apoptosis, we stained control and LACTB KD cells with Annexin-V and 7-AAD(*25*). LACTB KD cells display reduced apoptotic cell percentage compared to control upon staurosporine treatment (Fig. 1D and E, Fig. S1D-I). As an alternative apoptosis inducer, we treated cells with ABT-737(*26*) (Bcl-2 inhibitor) and S63845(*27*) (MCL-1 inhibitor) to activate the BAX pore(*28*, *29*). As with staurosporine, LACTB KD cells display increased cell viability upon this treatment (Fig. 1F, Fig. S 1 C).

Previous studies on LACTB-mediated tumour suppression showed that LACTB over-expression (OE) caused a decrease in cell proliferation and tumour growth(*14*, *15*, *17–19*). We therefore stably over-expressed LACTB in HeLa cells (Fig. 1G) and tested their response to apoptosis. LACTB-OE cells display increased sensitivity to cell death induced by staurosporine (Fig. 1H) or ABT-737/S63845 (Fig. 1I). Taken together, these results suggest that LACTB plays a role in apoptosis.

### LACTB is involved in release of mitochondrial proteins during apoptosis

Next, we investigated the mechanism by which LACTB enhances apoptosis. A previous study(*14*) suggested that LACTB functions by proteolytically degrading the phosphatidylserine decarboxylase (PISD) enzyme. However, we did not detect significant changes in PISD levels upon LACTB KD or OE (Fig. S2A,B), similar to findings by others(*15*, *16*, *30*). We therefore sought alternate mechanisms.

MOMP is often a committed step in apoptosis, leading to the release of mitochondrial proteins such as cytochrome c, SMAC/Diablo, AIF, and HTRA2/Omi into the cytoplasm(*1*, *31–33*). Using differential centrifugation to separate cytoplasm from mitochondria(*31*), we observed an increase in cytoplasmic levels of these mitochondrial proteins upon treatment with staurosporine or ABT-737/S63845 in control cells (Fig. 2A). LACTB KD causes a significant reduction in cytosolic release of these mitochondrial proteins induced by staurosporine (Fig. 2B) or ABT-737/S63845 (Fig. 2C). The overall levels of these proteins remains unchanged with either LACTB KD or OE (Fig. S2C,D).

**Figure 2:**
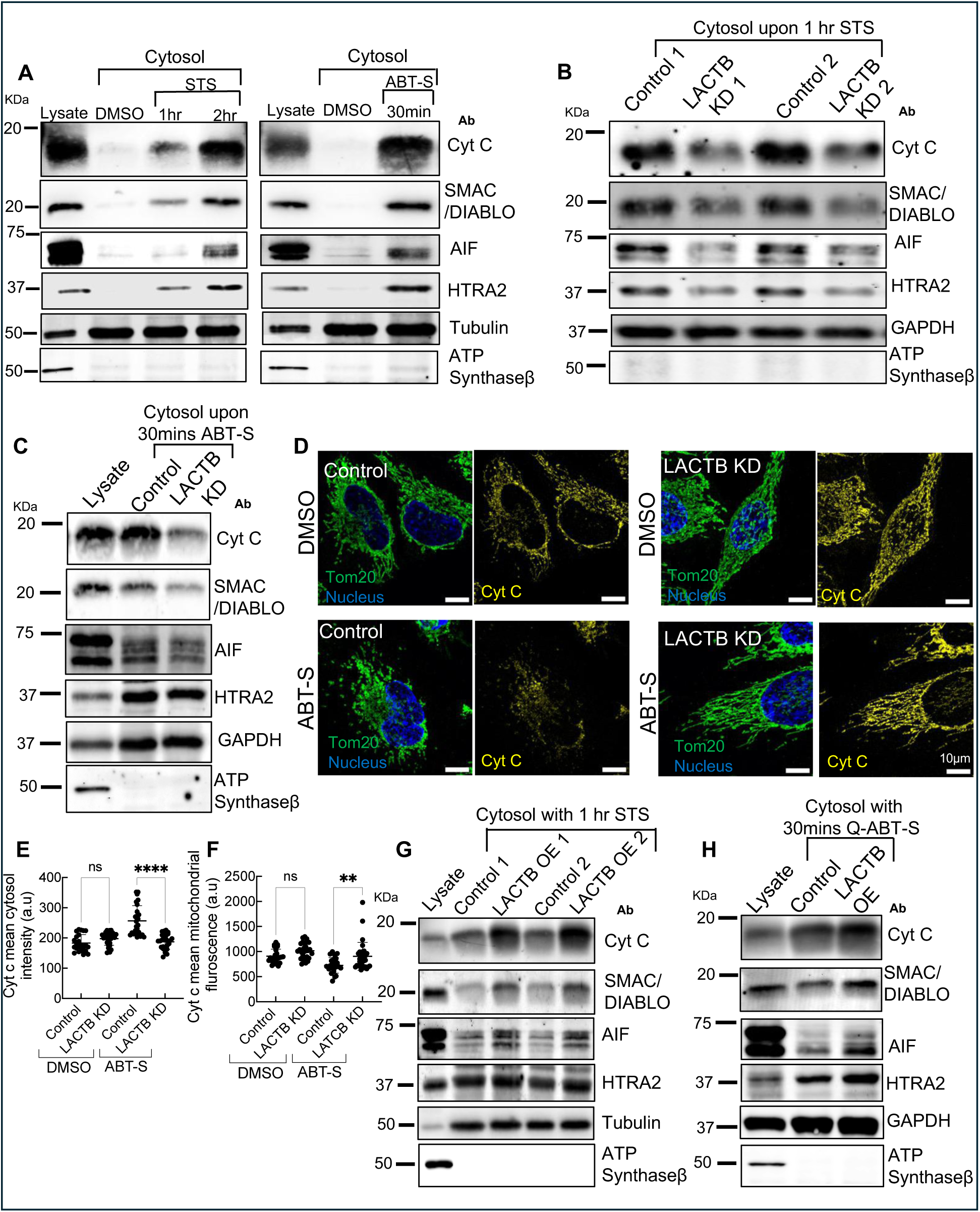
LACTB enhances mitochondrial release of cytochrome c and other factors upon apoptosis. (A) Western blot showing time-course analysis of mitochondrial apoptotic factor release into the cytosol following 1 μM STS or ABT-S treatment in HeLa cells. A total of 6 and 10 µg of protein was loaded per sample for STS and ABT-S treatment respectively. (B) Western blot showing mitochondrial apoptotic factor release into the cytosol after 1 μM STS treatment for 1 hour in control or LACTB KD HeLa cells. A total of 14 µg of protein was loaded per sample. (C) Western blot showing mitochondrial apoptotic factor release into the cytosol after ABT-S treatment for 30 minutes in control or LACTB KD HeLa cells. A total of 12 µg of protein was loaded per sample. (D) Fluorescence staining of cytochrome c, Tom20, and DNA (DAPI) in control and LACTB KD HeLa cells under DMSO and ABT-S treatment for 30 minutes. (**E**, **F**) Quantification of cytosolic cytochrome c levels (E) and mitochondrial cytochrome c retention (F) in from fluorescence staining experiments shown in panel d. N ≥ 23 cells; ****p < 0.0001, **p < 0.01 (one-way ANOVA). (G) Western blot showing mitochondrial apoptotic factor release into the cytosol after 1 μM STS treatment for 1 hour in control or LACTB OE HeLa cells. A total of 10 µg of protein was loaded per sample. (H) Western blot showing mitochondrial apoptotic factor release into the cytosol after ABT-S treatment for 30 minutes in control or LACTB OE HeLa cells. Cells treated with the pan-caspase inhibitor Q-VD-OPh (20 μM, 1-hour preincubation) prior to ABT-S treatment. A total of 12 µg of protein was loaded per sample.

Additionally, we used immunofluorescence to monitor cytochrome c release. In control HeLa cells, cytochrome c staining is concentrated in mitochondria, but largely shifts to cytosol after 30mins ABT-737/S63845 treatment (Fig. 2D). LACTB KD reduces cytochrome c translocation (Fig. 2D-F). We observed similar results in U2-OS cells with staurosporine treatment (Fig. S2E-G).

Finally, we tested the effect of LACTB OE on mitochondrial release of cytochrome c and other factors. LACTB OE leads to increased staurosporine-induced release of mitochondrial factors (Fig. 2G). To determine whether this effect is independent of caspase activity, we incubated cells with the pan-caspase inhibitor Q-VD-OPh prior to apoptosis induction(*34*). Even under these conditions, LACTB OE facilitates the ABT-737/S63845-induced release of cytochrome c and other mitochondrial proteins during apoptosis (Fig. 2H). In subsequent experiments, we employ caspase inhibition to prevent cell detachment while imaging. Taken together these results suggest that LACTB regulates release of mitochondrial proteins during apoptosis.

### LACTB is required for mitochondrial remodelling upon apoptosis

Next, we asked how LACTB might participate in release of mitochondrial factors during apoptosis. We first examined LACTB localization. An earlier study(*13*) suggested LACTB to be present in the mitochondrial intermembrane space. We re-examined LACTB localization by immunofluorescence using Airyscan microscopy in HeLa cells stably over-expressing un-tagged LACTB, relative to: an outer mitochondrial membrane marker (Tom20-GFP), and a mitochondrial matrix marker (Mito-GFP). LACTB staining is consistently found within the Tom20 signal (Fig. S3A-B), and surrounding the Mito-GFP signal (Fig. S3C-D). We also performed immunofluorescence staining for LACTB and ATP synthase beta subunit (on the matrix side of cristae). The LACTB and ATP synthase signals display strong colocalization (Fig. S3E, magnified insert 3F and line profile 3G), as supported by a positive correlation analysis (Fig. S3H). These results suggest that LACTB localizes to the IMM, and possibly enriches in cristae.

One possible mechanism for LACTB-mediated enhancement of mitochondrial protein release is through facilitating BAX/BAK channel assembly on the outer mitochondrial membrane(*35*). First, we checked BAX and BAK levels in control, LACTB KD and LACTB OE cells, and observed no significant differences (Fig. S4A). BAK is single-pass transmembrane protein constitutively present on the outer mitochondrial membrane (OMM), but BAX is recruited to the OMM from the cytosol upon apoptotic stimulation(*36*). To assess whether LACTB KD affects mitochondrial BAX recruitment, we immunostained for BAX under apoptotic and non-apoptotic conditions. Both control and LACTB KD cells display negligible mitochondrial BAX under DMSO treatment (Fig. S4B). Upon ABT-737/S63845 treatment, BAX punctae accumulate on mitochondria in both control and LACTB KD cells (Fig. S4C and D). Cell fractionation analysis further corroborates these findings (Fig. S4E). These results suggest that LACTB facilitates mitochondrial protein release without directly influencing BAX recruitment to mitochondria.

Another possibility is that LACTB mediates changes within mitochondria, which in turn facilitates mitochondrial protein release. The inner mitochondrial membrane undergoes extensive remodelling during apoptosis(*37*),(*9*). We examined mitochondrial morphology in control and LACTB KD HeLa cells, using Tom20 immunostaining and Airyscan microscopy. In the absence of apoptotic stimulus (DMSO control), mitochondria in LACTB KD cells are similar morphology to those in control KD cells (Fig. S5A). Upon treatment with staurosporine or ABT-737/S63845, control mitochondria change morphology substantially (Fig. 3A-C and Fig. S5B,C), with one change being an increase in mitochondrial width (Fig. S5D). In contrast, mitochondria in LACTB KD cells do not change noticeably in morphology (Fig. 3B,C and Fig. S5B-D). Quantification of mitochondrial width shows that control and LACTB KD cells display statistically indistinguishable mitochondrial widths in the absence of apoptotic stimulation (Fig. 3D). Upon staurosporine or ABT-737/S63845 treatment, mitochondria in control cells display an approximate 2-fold increase in width, while those in LACTB KD cells retain widths similar to DMSO treatment (Fig. 3D). We further tested these results by using ATP synthase as a marker for mitochondrial width. While the overall widths measured for ATP synthases are ∼ 100nm smaller than for Tom20 (which is expected, due to their relative locations), a similar trend occurs, with LACTB KD eliminating the width change induced by apoptotic stimulation (Fig. 3E, Fig. S6A-D).

**Figure 3.**
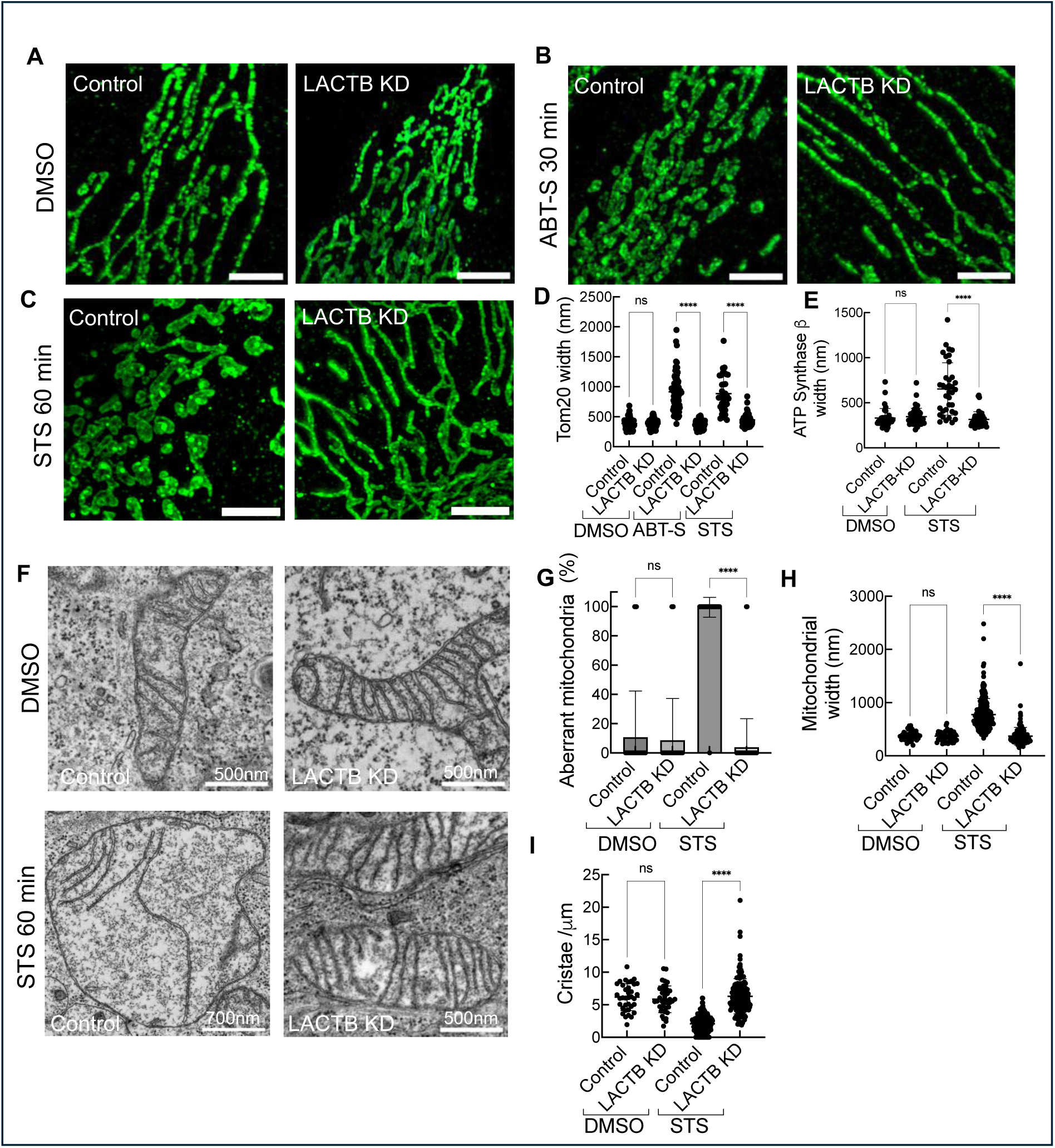
LACTB mediates mitochondrial remodelling upon apoptosis induction. (**A**-**C**) Immunofluorescence staining of Tom20 (green) in control siRNA and LACTB KD HeLa cells treated with DMSO (A, 1 hour), ABT-S (B, 30 min), or STS (C, 1 hour). Scale bar =5 mm. (**D**,**E**) Quantification of mitochondrial width changes under DMSO, ABT-S, or STS treatment (as in panels A-C) using Tom20 staining (D) or ATP synthase b staining (E). Widths determined by Gaussian fitting and full-width at half-maximum analysis. N = 2 independent experiments. N_mito_ (DMSO) ≥ 40; N_mito_ (ABT-S or STS) ≥ 75. (F) TEM micrographs of mitochondria in control and LACTB KD cells under DMSO or STS treatment for 1 hour. (G) Quantification of aberrant mitochondria from analysis of TEM micrographs of control and LACTB KD HeLa cells under DMSO or STS treatment for 1 hour. (H) Mitochondrial widths measured from TEM micrographs of control and LACTB KD HeLa cells under DMSO or STS treatment for 1 hour . (I) Quantification of cristae density (number per micron) from TEM micrographs of control and LACTB KD HeLa cells under DMSO or STS treatment for 1 hour. For panels (F–I), all quantifications were performed in a blinded manner. Error bars represent mean ± SD. Control DMSO_mito_ = 38; LACTB KD DMSO_mito_ = 46; Control STS_mito_ = 213; LACTB KD STS_mito_ = 153. N = 3 independent experiments. ****p < 0.0001; ns, not significant (one-way ANOVA).

To examine ultrastructural changes during apoptosis, we performed thin-section electron microscopy (EM) on control and LACTB KD cells. Consistent with the fluorescence microscopy results, mitochondrial widths are unchanged by LACTB KD under DMSO treatment. Upon staurosporine treatment, control cells display extensive mitochondrial swelling and large-scale morphological changes, whereas LACTB KD cells do not (Fig. 3F and G, Fig. S5E). Quantification of thin-section EM images reveals an approximate two-fold increase in mitochondrial width in control cells under staurosporine treatment (similar to our immunofluorescence results), while mitochondrial widths in LACTB KD cells do not change significantly (Fig. 3H). Additionally, there is a significant reduction in the number of cristae in control cell mitochondria upon apoptotic induction, a feature absent in LACTB KD cells (Fig. 3I). We also employed a recently developed fluorescent dye that stains cristae, PKmito Orange FX(*38*). Consistent with our EM and immuno-fluorescence data, control cell cristae exhibit extensive remodelling following apoptotic induction with staurosporine or ABT-737/S63845. In contrast, cristae in LACTB KD cells retain their structure, resembling those in untreated samples (Fig. S6E-I).

To investigate the kinetics of mitochondrial shape change in live cells during apoptosis, we transfected a matrix marker (Mito-dsRed) into control and LACTB KD cells. We also monitored cytochrome c release kinetics using a previously described cytochrome c-GFP construct(*23*, *39*). Prior to imaging, cells were treated with a pan-caspase inhibitor (Q-VD-OPh) for one hour, and then mounted on the microscope and stimulated with ABT-737/S63845. In control cells, cytochrome c release occurs at ∼ 28 minutes post-stimulation (Fig. 4A,B and E Supplementary Movie 1), followed by mitochondrial shape changes at ∼33 minutes (Fig. 4A, F; Supplementary Movie 2). In contrast, LACTB KD cells exhibit a significant delay in both cytochrome c release and mitochondrial remodelling (Fig. 4C-F; Supplementary Movie 3-4). Moreover, a subpopulation of LACTB KD cells displays neither cytochrome c release nor mitochondrial shape change over the 1-hr imaging period.

**Figure 4.**
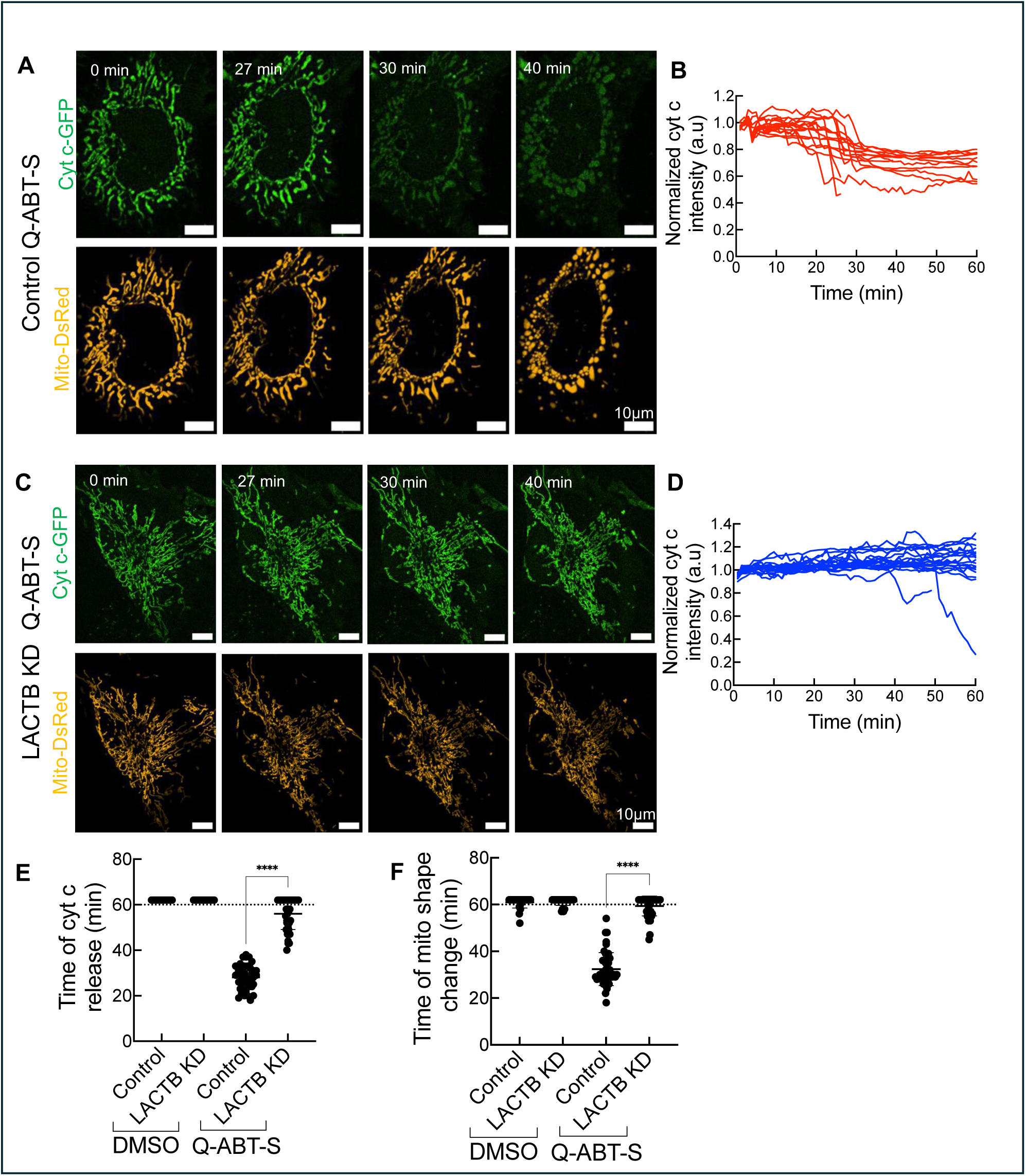
LACTB depletion slows cytochrome c release and mitochondrial remodelling kinetics upon apoptotic stimulation. (A) Live-cell imaging of cytochrome c release (Cyt c-GFP) and mitochondrial shape changes (Mito-DsRed) in HeLa cells treated with ABT-S in the presence of the pan-caspase inhibitor Q-VD-OPh (20 μM, 1-hour preincubation). (B) Quantification of cytochrome c release kinetics in control siRNA-treated cells treated with Q-ABT-S. N = 3 independent experiments, N_cells_= 16. (C) Live-cell imaging of cytochrome c release and mitochondrial shape changes in LACTB KD cells treated with Q-ABT-S and Q-VD-OPh (as in panel a). (D) Quantification of cytochrome c release kinetics in LACTB KD cells treated with Q-ABT-S. N = 3 independent experiments, N_cells_= 18. (E) Quantification showing the onset of cytochrome c release in control and LACTB KD cells under DMSO and Q-ABT-S treatment. N = 3 independent experiments, N_cells_= 39 (F) Quantification showing the onset of mitochondrial shape changes in control and LACTB KD cells under DMSO and ABT-S treatment. N = 3 independent experiments, N_cells_= 39 For panels (E–F), all quantifications were performed in a blinded manner. No cytochrome c release or mitochondrial shape changes were observed for data points above the dotted line. Error bars represent mean ± SD. N indicates the number of independent experiments. ****p < 0.0001; ns, not significant (one-way ANOVA).

Mitochondrial membrane remodelling can also be induced by CCCP, a potent mitochondrial depolarizer(*40*, *41*). We tested whether LACTB KD alters CCCP-induced mitochondrial remodelling. There is no difference in mitochondrial remodelling between control and LACTB KD cells upon CCCP treatment (Fig. S7A,B).

Taken together, these results demonstrate that LACTB plays a critical role in regulating mitochondrial morphology and cristae remodelling during apoptosis. The fact that LACTB KD does not alter remodelling caused by mitochondrial depolarization suggests that LACTB’s role is function-specific, rather than a general effect.

### LACTB-mediated mitochondrial remodeling is independent of Opa1 processing

The proteolytic processing of OPA1 has previously been implicated in IMM remodelling during apoptosis(*11*). To investigate whether the resistance to mitochondrial shape changes observed in LACTB KD cells was due to changes in OPA1 levels or processing, we examined OPA1 by western blotting. A slight increase in OPA1 levels occurs in LACTB KD cells compared to controls (Fig. S7C). However, OPA1 processing to the short form occurs normally in LACTB KD cells following treatment with Q-ABT-737/S63845 (Fig. S7C). We also tested OPA1 processing during staurosporine-induced apoptosis. Interestingly, no clear changes in OPA1 processing occur in either control or LACTB KD cells following one hour of staurosporine treatment (Fig. S7C), a time period sufficient to induce cytochrome c release and mitochondrial shape changes. We also tested the effect of Opa1 KD on cytochrome c-GFP release upon Q-ABT-737/S63845 treatment. Consistent with previous report(*39*), Opa1 KD does not significantly alter the kinetics of cytochrome c release (Fig. S7E-F). These results suggest that LACTB mediates mitochondrial remodelling in a manner distinct from Opa1.

### LACTB binds and remodels cardiolipin-containing curved membranes

We have shown that LACTB: 1) acts in apoptosis by facilitating release of cytochrome c and other factors from mitochondria, 2) co-localizes with ATP synthase, and 3) plays a role in mitochondrial morphology changes during apoptosis. LACTB has previously been reported to bind cardiolipin-containing membranes(*42*). To investigate whether LACTB actively participates in membrane remodeling, we tested the binding of purified LACTB-GFP (Fig. S8A, B) to supported lipid membranes, using a system in which membrane nanotubes are pulled from a supported lipid bilayer (SLB) (*43*, *44*). These nanotubes mimic cristae dimensions (20-60 nm diameter) (Fig. S8C). This assay offers the ability to monitor a range of membrane curvatures simultaneously, from the planar SLB to nanotubes of varying diameter.

Efficient membrane binding by LACTB-GFP requires ∼25 mol% cardiolipin (Fig. 5A; Fig. S8D). This requirement is not solely due to cardiolipin’s negative charge, as equivalent charge density of phosphatidylserine is insufficient for binding (Fig. 5B; Fig. S8D). Interestingly, LACTB-GFP binding occurs mainly on nanotubes as well as at the fringes of the SLB, suggesting that LACTB exhibits a strong preference for curved membranes (Fig. 5C, D).

**Figure 5:**
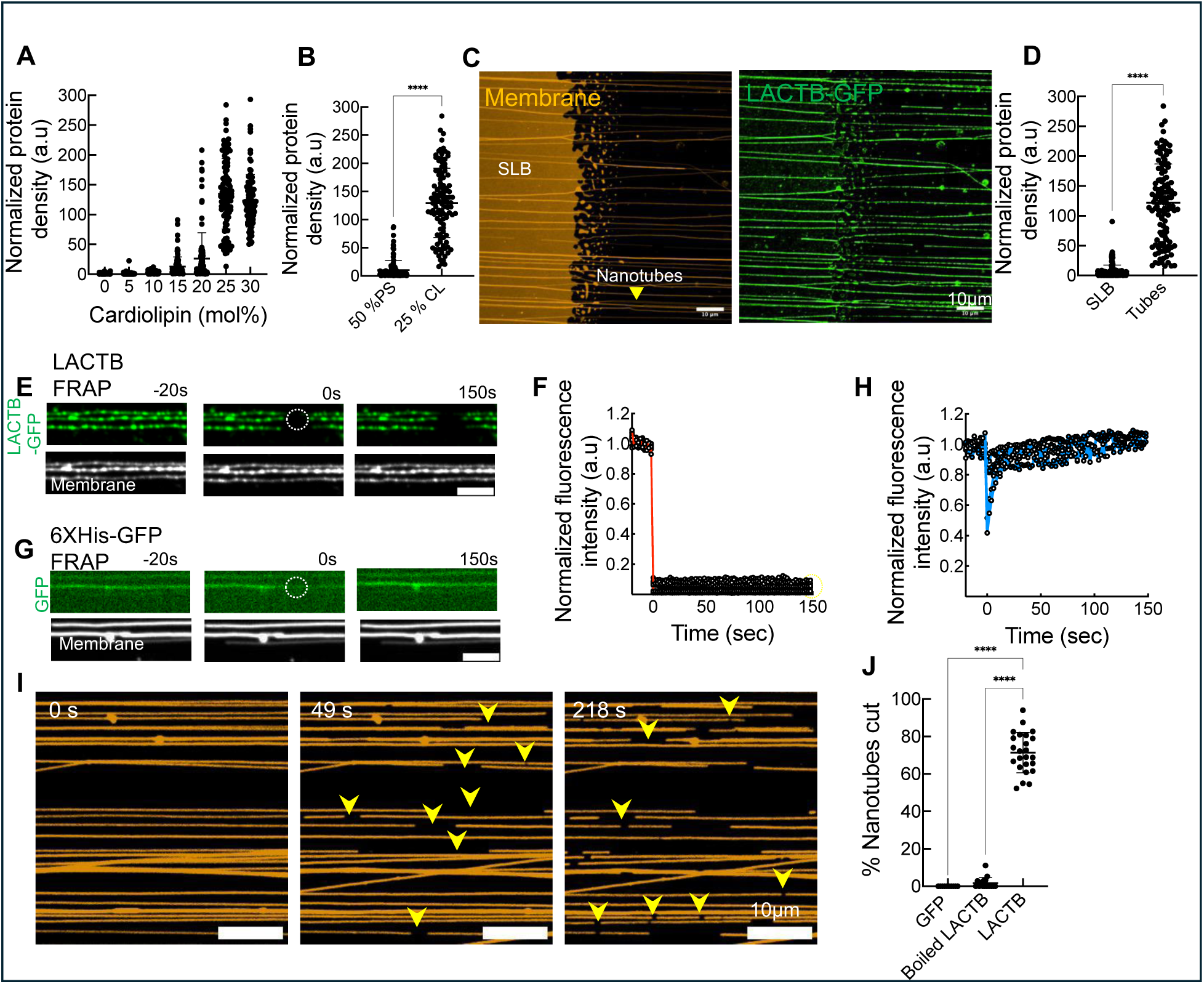
LACTB preferentially binds and remodels curved membranes containing cardiolipin. (A) Binding of 1μM LACTB-GFP to membrane nanotubes in response to increasing cardiolipin concentration. Lipid mixtures contained DOPC and varying levels of cardiolipin, with 0.5 mol% RhPE for visualization. N_tubes_ ≥ 80. (B) Quantification of 1μM LACTB-GFP binding to nanotubes containing either 25 mol% cardiolipin or 50 mol% phosphatidylserine. N_tubes_ ≥ 103. (**C**, **D**) Representative micrographs and quantification of 1μM LACTB-GFP (green) binding on nanotubes compared to supported lipid bilayers (SLBs). Lipid composition: DOPC:CL:RhPE (74.5:25:0.5 mol%). N=3; N_SLB_ = 155; N_tubes_ = 119. (**E**,**F**) Representative images (E) and fluorescence recovery curves (F) showing FRAP kinetics of LACTB-GFP on nanotubes containing 25 mol% cardiolipin. N = 2; N_tubes_ = 15. Scale bar = 5μm. (**G**,**H**) Representative images (G) and fluorescence recovery curves (H) showing the FRAP kinetics of 6XHis-GFP alone on membrane nanotubes containing 5 mol% Ni-NTA. N = 1; N_tubes_ = 6 Scale bar = 5μm. (I) Stills from a time-lapse movie showing the membrane marker RhPE upon LACTB (1 μM) addition to nanotubes containing 25 mol% cardiolipin for the indicated times. Yellow arrowheads indicate sites of fission on the nanotubes. (J) Quantification of nanotube fission after incubation with 1μM LACTB, boiled LACTB, or GFP for 10 minutes. Each data point represents an entire field containing 10–15 nanotubes. N = 3 independent experiments, N_tubes_≥ 104. Error bars represent mean ± SD. ****p<0.0001; ns, not significant (Mann–Whitney test).

Dynamic monitoring of LACTB-GFP binding to nanotubes reveals punctate assembly, with punctae nucleating at specific sites on the tubes (Fig. S8E). The assembly process follows a half-time (t₁/₂) of 52.9 ± 3 seconds (Fig. S8F). Once formed, these ‘scaffolds’ do not exchange with free LACTB-GFP in solution, as demonstrated by fluorescence recovery after photobleaching (FRAP) analysis (Fig. 5E,F). This behaviour contrasts with 6XHistidine-tagged GFP binding to nanotubes containing Ni-NTA-lipid, which displays uniform binding and rapid recovery after FRAP (Fig. S8E and Fig. 5G,H).

LACTB binding also induces nanotube fission, observed in real time (Fig. 5I; Supplementary Movie 5; Fig. S9A). These fission events require properly folded LACTB, as neither boiled LACTB nor soluble GFP cause nanotube fission (Fig. 5J; Fig. S9B, C). Montages of nanotubes undergoing fission reveals that a distinct drop in fluorescence intensity occurs at the fission site prior to fission, suggesting that LACTB induces membrane constriction *en route* to fission (Fig. S9D–F). Using kymograph analysis, we captured membrane intermediates preceding fission in some instances (Fig. S9G, H) but not in others (Fig. S9I). Together, these findings demonstrate that LACTB preferentially binds curved membrane surfaces in a punctate manner, and is capable of inducing membrane constriction as well as fission.

## Discussion

The mechanisms underlying IMM remodelling during apoptosis—as well as their role in cell death—remain controversial(*1*, *8*, *45*, *46*). Previous work has shown that the IMM dynamin Opa1 might play an inhibitory role in this process by maintaining cristae architecture, thereby delaying cytochrome c release(*11*). Several mechanisms might relieve this Opa1-mediated inhibition, including BH3-only proteins(*9–12*), the mitochondrial fission machinery(*47*, *48*), and proteolytic cleavage by the IMM protease OMA1(*49*, *50*). Conversely, the IMM protease PARL has been reported to cleave Opa1 in a manner that enhances its anti-apoptotic activity(*9*, *11*), although another study implicates PARL as pro-apoptotic via a different mechanism(*51*). Moreover, it remains unclear whether IMM remodelling is essential for apoptosis, as one study found that such remodelling occurs after cytochrome c release during etoposide-induced apoptosis(*37*).

Our work reveals three fundamental findings: 1) the IMS protein LACTB enhances apoptosis through promoting mitochondrial protein release, 2) LACTB also is required for mitochondrial remodelling during apoptosis, and 3) purified LACTB can directly remodel phospholipid membrane nanotubes. Importantly, LACTB function is not directly related to mitochondrial BAX recruitment, but to mitochondrial remodelling. The fact that LACTB KD inhibits apoptosis-induced mitochondrial remodelling, but not remodelling caused by CCCP, suggests that LACTB’s function is purpose-specific rather than as a general remodelling factor. In addition, altering LACTB levels by KD do not change Opa1 processing, suggesting that LACTB’s effects are independent of Opa1-induced mitochondrial remodelling. Our biophysical studies reveal that LACTB preferentially binds and remodels curved membranes enriched in cardiolipin, suggesting a direct role in mitochondrial membrane remodelling during apoptosis. These findings establish LACTB as a unique membrane remodelling factor in mitochondrial dynamics during apoptosis, distinct from mitochondrial dynamin proteins like Opa1, Drp1, and mitofusins(*44*).

Our live-cell assays indicate that apoptotic cytochrome c release precedes overt changes in mitochondrial morphology, consistent with previous findings(*37*). This observation raises the question of whether these two processes are functionally linked. Notably, LACTB KD strongly suppresses both cytochrome c release and mitochondrial morphological alterations, suggesting a potential relationship. One possibility is that subtle morphological changes necessary for cytochrome c release from cristae are below the detection threshold of our imaging system, and the larger-scale changes we observe occur after cytochrome c release. Alternatively, both events may depend on LACTB yet be mechanistically independent.

While one potential mechanism for LACTB-mediated IMM remodelling is its direct function as a membrane remodelling protein, another possibility is that LACTB stimulates membrane remodelling through other proteins. A previously proposed mechanism for LACTB function is through proteolysis of the PISD protein, an enzyme which converts phosphatidylserine to phosphatidylethanolamine(*14*). While we show that PISD processing is not altered by LACTB KD or OE, other proteins might be LACTB substrates during apoptosis, with one candidate being PLA2G6(*30*). As another possibility, LACTB could mediate contact between inner and outer mitochondrial membranes, facilitating cardiolipin transfer to the outer membrane—a process critical for BAX activation(*52–54*). Future studies will investigate the molecular mechanism of LACTB’s apoptotic role in more detail. It will also be interesting to determine whether LACTB plays a role in mitochondrially-derived vesicle/mitochondrially-derived compartment assembly(*55*, *56*) or in mitochondrial nucleic acid release upstream of inflammatory activation through cGAS-STING(*57*, *58*) and other pathways.

Our identification of a role for LACTB in mitochondrial protein release does not rule out other tumour-suppressing functions. In fact, it is possible that LACTB is itself released from mitochondria and plays additional roles through its previously reported interactions with p53(*15*), PP1A(*16*), and components of the PI3K/AKT signalling pathway(*20*). Further studies will investigate these potential functions, as well as the contributions of its enzymatic activity and polymerization ability.

Our work establishes LACTB as a critical regulator of mitochondrial morphology during apoptosis. By identifying its curvature-specific membrane remodelling activity and its distinct role in apoptotic signalling, we provide a foundation for exploring LACTB’s therapeutic potential, particularly in diseases where mitochondrial dynamics and apoptosis are dysregulated.

## Materials and Methods

### Cell culture

Human cervical cancer (HeLa cells), osteosarcoma (U2-OS) cells, and murine melanoma (B16-F10) cells were obtained from the American Type Culture Collection (ATCC). Cells were cultured in medium Dulbecco’s Modified Eagle Medium (DMEM; Corning, 10-013-CV) supplemented with 10% fetal bovine serum (FBS; Sigma-Aldrich, F4135) at 37°C in a humidified atmosphere containing 5% CO2. Mycoplasma contamination was monitored every three months using the MycoStrip® Mycoplasma Detection Kit (InvivoGen, rep-mys-50). All lines were maintained for no more than 30 passages.

### siRNA knockdown

Oligonucleotides for siRNA-mediated silencing were synthesized by Integrated DNA Technologies (IDT). For human LACTB, sequence targeting Exon 5 (3′ UTR) was 5′-ACUUGGAUAUGCUGACGACUGUGCA-3′. For mouse LACTB, sequences targeting CDS Exon 3 was 5′-AAAAGGUUUCUGUCACAACAAGATT-3′. Human OPA1 was targeted using sequence to Exon 19–22: 5′-CCACAGUGGAUAUCAAGCUUAAACA-3′. Negative control siRNA sequence was 5′-CGUUAAUCGCGUAUAAUACGCGUAU-3′.

Cells were seeded at 7.5 × 10^4^ per well in 6-well dishes and cultured overnight in 2 mL medium. The following day, transfection mixes were prepared by incubating 63 pmol of each siRNA with 50 µL Opti-MEM (Gibco, 31985-070) and 2 µL RNAiMAX (Thermo Fisher, 13778) with 100 µL Opti-MEM separately for 10 minutes. The two solutions were then combined, and incubated for an additional 20 minutes. Medium was replaced with 1 mL fresh medium, and 150 µL of the transfection mix was added dropwise. After 10 hours, medium was replaced with 2 mL medium. Transfections were repeated 48 hours later. At 72 hours after the initial transfection, cells were re-plated at seeding density of 2 × 10^5^ cells onto fibronectin-coated (Sigma, F1141-1MG) MatTek dishes (MatTek, P35G-1.5-14-C) and processed for fixation or live cells microscopy.

### Plasmids and transfections

GFP–Mito was purchased from Clontech (pAcGFP1-Mito, 632432) and contains the mitochondrial targeting sequence derived from the precursor of subunit VIII of human cytochrome c oxidase. Mito–DsRed was previously described(*59*). Tom20–GFP was previously described(*40*). Cytochrome C–GFP was obtained from Douglas Green (Addgene plasmid #41182). The Super PiggyBac Transposase was purchased from System Biosciences LLC (PB210PA-1), and the PiggyBac Dual Promoter (PB513B-1) was purchased from Lifescience Market (empty plasmid used as control for overexpression). The human LACTB coding sequence was PCR-amplified from a sequence-verified cDNA clone (Accession: BC067288; Clone ID: 30366453; Dharmacon, MHS6278-202804693), and inserted into the PiggyBac Dual Promoter vector using XbaI and NotI restriction sites, and into the eGFP–N1 mammalian expression vector (Clonetech) using XhoI and KpnI restriction sites. Plasmid transfections were initiated by seeding cells at 4 × 10⁵ cells per well in 35-mm dishes approximately 16 hours prior to transfection. Transfections were carried out in OPTI-MEM medium (Gibco, 31985062) using Lipofectamine 2000 (Invitrogen, 11668) according to the manufacturer’s protocol. Post-transfection, cells were trypsinized and re-plated onto fibronectin coated glass-bottom dishes (MatTek Corporation, P35G-1.5-14-C) at a density of ∼1 × 10⁵ cells per well. Imaging was performed approximately 16–24 hours after transfection. For preparation of stable cell lines by the Piggybac system, transfections were performed following the manufacturer’s protocol (System Biosciences LLC), and puromycin selection (2 µg/mL) was applied at 24 hr post-transfection for 3–5 days to establish stably transfected cell lines.

### Antibodies

Details of all antibodies used in this study, including their sources, catalog numbers, and dilutions, are provided in Supplementary Table 1

### Drug treatments

Staurosporine (Research Product International, S63500-0.0001, referred to as STS in figures) was used at 1 mM. The combination of ABT-737 (10 µM, ApexBio, A8193) and S63845 (2 µM, ApexBio, A8737) was collectively referred to as ABT-S in figures. CCCP (20 µM for 20 mins, Sigma-Aldrich, C2759). For experiments involving the pan-caspase inhibitor Q-VD-OPh (20 µM) (ApexBio, A1901), cells were preincubated for 1 hour prior to ABT-S treatment (referred to as Q-ABT-S in figures). All treatments were performed in prewarmed medium and incubated at 37°C with 5% CO₂ for the indicated durations.

### Sulforhodamine B (SRB) assay

Assay was performed as per reported protocol(*24*). Briefly, 1 × 10⁴ cells were plated per well (100 mL) in a 96-well plate and incubated overnight. The next day, 100 µL of medium containing either the drug of interest or DMSO was added to each well. At the end of the treatment, media was aspirated; cells were fixed by adding 100 µL 10% trichloroacetic acid (TCA; Sigma-Aldrich, T6399) per well and incubating for 30 minutes at 4°C. The TCA was then removed, and the wells were washed five times with MilliQ water using a multichannel pipette. Plates were air-dried completely before staining each well with 100 µL of 0.4% SRB (Sigma-Aldrich, 230162) in 1% acetic acid for 10 minutes at room temperature. Excess dye was removed by washing the wells five times with 1% acetic acid, followed by air drying. The bound dye was solubilized by adding 100 µL of 10 mM Tris-HCl (pH 8.0) to each well and gently shaking for 5 minutes. Absorbance was measured at 520 nm using a M-1000 plate reader (Tecan Inc) through i-control software (version1.11). Each data point in the SRB assay graph represents an individual well (i.e., technical replicates)

### Annexin V staining

Cells were harvested using 0.05% Trypsin, washed with 1X PBS (Corning, 21-040-CM), counted, and normalized to a concentration of 1 × 10⁶ cells/mL. Apoptosis and cell death markers were stained using PE Annexin V Apoptosis Detection Kit with 7-AAD (BioLegend, 640934) following the manufacturer’s protocol. Cells were washed twice with staining buffer (PBS with 2% newborn calf serum (Gibco, 26010074)) and resuspended in 50 µL of Annexin V Binding Buffer per sample. 2 µL of PE Annexin V and 2 µL of 7-AAD Viability Staining Solution were added. Cells were gently mixed and incubated for 15 min. at room temperature in the dark. Following incubation, 150 µL of Annexin V Binding Buffer was added to achieve a final volume of 200 µL. Flow cytometry was performed immediately using a CytoFLEX S flow cytometer (Beckman Coulter), and data were analyzed with FlowJo v10 software (BD Biosciences).

### Cytosol isolation

Protocol adapted from Bossy-Wetzel et al(*31*). Cells were trypsinized and resuspended in DMEM supplemented with 10% FBS to neutralize the trypsin. The cells were then washed with 1× PBS to remove excess medium and pelleted by centrifugation at 200 × g for 10 minutes at 4°C. The resulting pellet was resuspended in 500 µL of extraction buffer (220 mM mannitol, 68 mM sucrose, 20 mM HEPES, pH 7.4, 50 mM KCl, 2 mM MgCl₂, 1 mM DTT, and protease inhibitors) and incubated on ice for 30 minutes. Subsequently, the cells were homogenized for 20 strokes using a prechilled Dounce homogenizer (Wheaton Dura-Grind, 357574) and centrifuged at 14,000 × g for 15 minutes at 4°C. The pellet (nuclear, ER and mitochondrial fraction) and the supernatant (cytosolic fraction) were separated. Protein concentration in the cytosolic fraction was measured using the BCA assay (Thermo Fisher 23225) and and samples were prepared with 4X SDS–PAGE sample buffer diluted to 1X.

### Western blot

For cell extracts, lysate preparation involved washing confluent cells grown in 35-mm dishes three times with 1XPBS and lysing them in ∼400 µL of 1× DB buffer, consisting of 50 mM Tris-HCl (pH 6.8), 2 mM EDTA, 20% glycerol, 0.8% SDS, 0.02% bromophenol blue, 1,000 mM NaCl, and 4 M urea. Lysates were heated at 95°C for 10 minutes, and genomic DNA was sheared by sonicating for 10-minute in bath sonicator. For cytosolic and mitochondrial fractions, samples were prepared as described above. Proteins were resolved using SDS–PAGE and transferred onto polyvinylidene difluoride (PVDF) membranes (Millipore). Blocking was performed in TBS-T buffer (20 mM Tris-HCl, pH 7.6, 136 mM NaCl, 0.1% Tween-20) containing 3% bovine serum albumin (BSA) for 1 hour at room temperature, followed by overnight incubation with primary antibodies at 4°C. After three washes with TBS-T, membranes were incubated for 1 hour at room temperature with horseradish peroxidase (HRP)-conjugated secondary antibodies. Excess secondary antibodies were removed by three additional TBS-T washes. Chemiluminescence signals were detected using ECL Prime Western Blotting Detection Reagent (Cytiva Amersham, 45-002-40) and recorded with an ECL Chemocam imager (SYNGENE G:BOX Chemi XRQ). For fluorescence-based detection, membranes were incubated with IRDye 680 goat anti-mouse or IRDye 800CW goat anti-rabbit secondary antibodies for 1 hour at room temperature, and signals were visualized using the LI-COR Odyssey CLx imaging system. Information regarding primary and secondary antibodies is provided in Supplementary Table 1.

### Immunofluorescence staining

Cells were plated onto fibronectin-coated MatTek dishes 16 hours prior to fixation and staining. Treatments were performed at 37°C and 5% CO₂, after which cells were washed twice with PBS and fixed in either 1% glutaraldehyde (EMS, 16020) for 10 minutes or 4% prewarmed paraformaldehyde (EMS, 15170) prepared in PBS for 20 minutes. Fixed cells were permeabilized using 0.1% Triton X-100 in PBS for 10 minutes and blocked in PBS containing 10% calf serum for 1 hour. Cells were stained with the appropriate primary antibody (Supplementary Table 1) diluted in PBS containing 1% calf serum for 1.5 hours, followed by three PBS washes and incubation with the corresponding secondary antibodies (Supplementary Table 1) and DAPI (4’,6-diamidino-2-phenylindole (Calbiochem, 268298) for nuclear staining in PBS containing 1% calf serum for 1 hour. After washing in PBS, imaging was conducted in PBS using either Dragonfly spinning disk or Airyscan.

### Airyscan microscopy

Airyscan imaging was conducted using an LSM 880 confocal microscope equipped with a 100×/1.4 NA Apochromat oil objective and Airyscan detectors (Carl Zeiss Microscopy). Imaging was performed with a 488 nm laser and a Band Pass (BP) 420–480/BP 495–620 filter for Fluoresceine-labelled secondary antibodies, and a 561 nm laser with a BP 495–550/Long Pass (LP) 570 filter for Texas Red-labelled secondary antibodies. Z-stack images were acquired from the basal to apical regions with a step size of 0.2 µm. Raw image data were processed using the Airyscan processing feature in Zen Black software (Carl Zeiss, version 2.3). Maximum intensity projections were generated from z-stacks, and background subtraction was performed in ImageJ Fiji software using the rolling ball algorithm (radius: 20).

### Live cell microscopy using spinning disk confocal microscope

Live-cell imaging was performed in Dulbecco’s Modified Eagle Medium (DMEM; Corning, 10-013-CV) supplemented with 10% fetal bovine serum (FBS; Sigma-Aldrich, F4135). Approximately 2 × 10⁵ cells were plated onto fibronectin-coated MatTek dishes 16 hours before imaging. The medium was pre-equilibrated at 37°C and 5% CO₂ prior to use. Cells were preincubated for an hour prior to imaging with the pan-caspase inhibitor Q-VD-OPh (20 µM) (ApexBio, A1901) in DMEM media supplemented with 10% FBS.

Imaging was conducted using a Dragonfly 302 spinning disk confocal system (Andor Technology) on a Nikon Ti-E microscope base. The system was equipped with an iXon Ultra 888 EMCCD camera, a Zyla 4.2 Mpixel sCMOS camera, and a Tokai Hit stage-top incubator maintained at 37°C. Illumination was provided by a solid-state 405 smart diode laser (100 mW), a solid-state 560 OPSL smart laser (50 mW), and a solid-state 637 OPSL smart laser (140 mW). For drug treatment experiments, the compound was added at the start of the fourth image frame (∼4 minutes after imaging began; time interval set at 1 minute), and imaging continued for a total duration of 1 hour.

The CFI Plan Apochromat Lambda 100×/1.45 NA oil objective (Nikon, MRD01905) was used for live-cell drug treatment assays as well as fixed cell imaging. The CFI Plan Apochromat 60×/1.4 NA oil immersion objective was used for imaging during BAX localization experiments. Image acquisition was carried out using Fusion software (Andor Technology, version 2.0.0.15).

### PKmito Orange FX Staining

PKmito Orange FX (Cytoskeleton, Inc., CY-SC054) staining was conducted following the manufacturer’s protocol. Briefly, control and LACTB KD cells were seeded at an equal density (2 × 10⁵ cells) onto fibronectin-coated MatTek dishes 16 hours prior to imaging. On the day of imaging, cells were preincubated with medium containing 500 nM PKmito Orange FX at 37°C in a humidified atmosphere with 5% CO₂ for 1 hour. Drug treatments were carried out in the presence of 500 nM PKmito Orange FX for the indicated durations.

Following treatment, cells were washed three times with PBS and fixed with prewarmed 2.5% glutaraldehyde for a minimum of 30 minutes. The cells were then washed three more times with PBS, and DNA staining was performed by adding 0.1 mg/L (w/v) DAPI to the first wash to label the nucleus. Imaging was performed using Airyscan microscope.

### Thin section EM

Control and LACTB KD cells were seeded at a density of 3 × 10⁵ cells/mL onto fibronectin-coated MatTek dishes 16 hours prior to fixation. On the day of fixation, cells were treated with staurosporine (STS) or DMSO (control) and incubated for 1 hour. Fresh fixative containing 0.1 M sodium cacodylate buffer, 2% glutaraldehyde, and 3.2% paraformaldehyde in Milli-Q water was prepared within an hour before use and warmed to 37°C in a tissue culture bath. Media was aspirated from the dish, and 2 mL of warm fixative was added immediately without prior washing. After 15 minutes, the fixative was replaced with 1.5 mL of fresh fixative from the same batch, and cells were incubated at room temperature for 1 hour. The dishes were sealed with parafilm and stored in an airtight container for subsequent processing. Post-fixation was carried out with 1% osmium tetroxide in 0.1 M sodium cacodylate buffer (pH 7.2) for 1 hour on ice in the dark, followed by rinsing twice with 0.1 M sodium cacodylate buffer for 10 minutes each at room temperature. Samples were incubated overnight in 2% aqueous uranyl acetate at room temperature in the dark, dehydrated through a graded ethanol series (50%, 70%, 95%, and 2 × 100%), and embedded in LX-112 resin (Ladd Research) using standard protocol(*60*). Sections (50 nm) cut parallel to the cellular monolayer were obtained using a Leica Ultracut 7 ultramicrotome, mounted on carbon/formvar-coated 300-mesh copper grids and stained with uranyl acetate and lead citrate. TEM micrographs were obtained using a Helios CX-5 electron microscope equipped with a STEM3+ detector at 20kV in high resolution immersion mode, and analysed using FIJI software, with blinded analysis.

### Protein expression and purification

2 × 10⁶ Expi293F™ cells (Life Technologies, A14527) were cultured in 1 liter of Expi293 expression medium (Life Technologies, A1435101) and transfected with 1 mg of DNA and 3 mg of sterile 25 kDa linear polyethyleneimine, prepared in Opti-MEM™ reduced-serum medium (Life Technologies, 31985070). The cells were incubated at 37°C with 8% CO₂ and shaking at 125 rpm for three days, harvested by centrifugation at 300 × g for 15 minutes at 4°C, and resuspended in 1× PBS.

All the subsequent steps were performed at 4°C or on ice. The cells were resuspended in 50 mL of lysis buffer (20 mM HEPES buffer, pH 7.4, 500 mM NaCl, 5 mM EDTA, 1 mM DTT, 2 μg/mL leupeptin, 10 μg/mL aprotinin, 2 μg/mL pepstatin A, 1 μg/mL calpeptin, 1 μg/mL calpain inhibitor I, 1 mM benzamidine, and 1:1,000 dilution of universal nuclease (Thermo Fisher, 88702)) per 5 mL cell pellet. The cells were lysed using a high-pressure homogenizer (M-110L; Microfluidics), and 1% Triton X-100 was added after solubilization. Cell debris was removed by centrifugation at 40,000 rpm (185,000xg) for 30 minutes using a Ti45 rotor (Beckman). The clarified supernatant was loaded onto Strep-Tactin Superflow resin (IBA, 2-1206-025) pre-equilibrated with lysis buffer, and washed with 30 column volumes wash buffer (10 mM HEPES, pH 7.4, 150 mM KCl, 1 mM EDTA, 1 mM DTT). The strep-tagged GFP fusion protein was eluted using strep elution buffer (10 mM HEPES, pH 7.4, 150 mM KCl, 1 mM DTT, and 2.5 mM desthiobiotin).

### Membrane nanotube assay

Lipid nanotubes were made as previously described(*43*, *44*), with all lipids sourced from Avanti Polar Lipids. Briefly, 2 μL of a 1 mM lipid stock (in chloroform) was deposited onto glass coverslips passivated with covalently attached PEG8000 (Sigma-Aldrich, P2139). The coverslips were vacuum-dried and assembled into a flow cell (FCS2, Bioptechs, PA). Lipids were hydrated by flowing 20 mM HEPES buffer, pH 7.4, containing 150 mM KCl. High flow rates of the same buffer facilitated the formation of a supported lipid bilayer (SLB) at the lipid source and generated membrane nanotubes downstream. The flow was then halted, allowing the nanotubes to settle and adhere to defects on the glass surface. Reaction mixtures were introduced onto the nanotubes at flow rates 10 times slower than those used during nanotube preparation. The SLB formed during this process served as an in situ calibration standard for estimating nanotube dimensions(*43*, *44*). Experiments involving LACTB-GFP on the nanotubes were conducted at a total protein concentration of 1 μM in binding buffer (20 mM HEPES buffer, pH 7.4, containing 150 mM KCl and 1 mM MgCl₂). Temperature was maintained at 37°C using a temperature controller from Bioptechs, PA.

### Fluorescence recovery after photobleaching (FRAP)

FRAP was performed on nanotubes following LACTB-GFP/6×His recruitment using a Dragonfly spinning-disk confocal microscope operated by Andor iQ3 software. Bleaching was achieved with a solid-state 405 smart diode laser (100 mW) set at 40% output power, delivering a 500 ms exposure to a fixed area of 2 µm². Images were captured in both the 488 nm (protein) and 561 nm (membrane) channels before bleaching and 1 second post-bleach. To minimize photobleaching during acquisition, imaging parameters were standardized, and the mean post-bleach intensity was normalized to the mean pre-bleach intensity.

### LACTB binding assay

LACTB-GFP (1 μM) was incubated with nanotubes of the respective membrane composition for 10 minutes, followed by washing with 400 μL of binding buffer. The nanotubes were subsequently imaged. Protein density was calculated by dividing the mean protein intensity by the mean membrane intensity and multiplying by 100, with the area held constant for both nanotubes and SLB.

### Image and statistical analysis

All graphical representations and statistical analyses were performed using GraphPad Prism (version 10.4.1). Statistical significance was determined using one-way ANOVA or the Mann– Whitney test, as appropriate. The number of independent experiments, number of cells or mitochondria analysed, and the statistical tests performed are provided in the respective figure legends. Image analysis was conducted using ImageJ Fiji (version 2.14.0/1.54f, National Institutes of Health). TEM micrographs were manually analysed using ImageJ after blinding. Mitochondria exhibiting swollen morphology and expanded cristae were classified as aberrant. Mitochondrial width was measured by drawing a line across the outer mitochondrial membrane from one side to the other, while cristae density was determined by counting the number of cristae within a mitochondrion and dividing by total mitochondrial length (µm). The number of experimental replicates for each assay is specified in the corresponding figure legends.

## Acknowledgments

We thank J. Delgado, A. Ferreira Verissimo, E. Shipman, C. Shoemaker, P. Sapoisto, and Z. Svindrych for their valuable assistance during these investigations. We also acknowledge M. Lee and Z. She for their help in blinding images and D. Andhare for critical feedback on the manuscript. We also thank Dartmouth Imaging Facility for Airy Scan microscopy and Dartmouth Electron Microscopy Facility for TEM imagining.

## Funding

This work was supported by

National Institutes of Health (NIH) R35 GM122545 (H.N.H)

National Institutes of Health (NIH) R01 DK088826 (H.N.H)

National Institutes of Health (NIH) R01 CA257954 (E.J.U.)

National Institutes of Health (NIH) R01 HL155824 (R.V.S.)

National Institutes of Health (NIH) P20 GM113132 (Dartmouth BioMT)

5P30CA023108 to the Dartmouth Cancer Centre (RRID:SCR_025077)

### Author contributions

S.C.K. performed and analyzed all cell biology, biochemical, and light microscopy experiments. T.K. carried out Annexin V staining and flow cytometry. R.V.S. prepared thin-section samples and conducted TEM imaging. H.N.H. and E.J.U. provided project supervision. S.C.K. and H.N.H. wrote the manuscript.

### Data and materials availability

All data supporting the conclusions of this study are available in the main text or Supplementary Figures 1–9. Additional data/materials supporting the findings of this study can be obtained from the corresponding author upon reasonable request.

## List of Supplementary Materials

Materials and Methods

Supplementary Text

Figs. S1 to S9 Table S1

Movies S1 to S5

## Supplementary Materials

**Figure S1.**
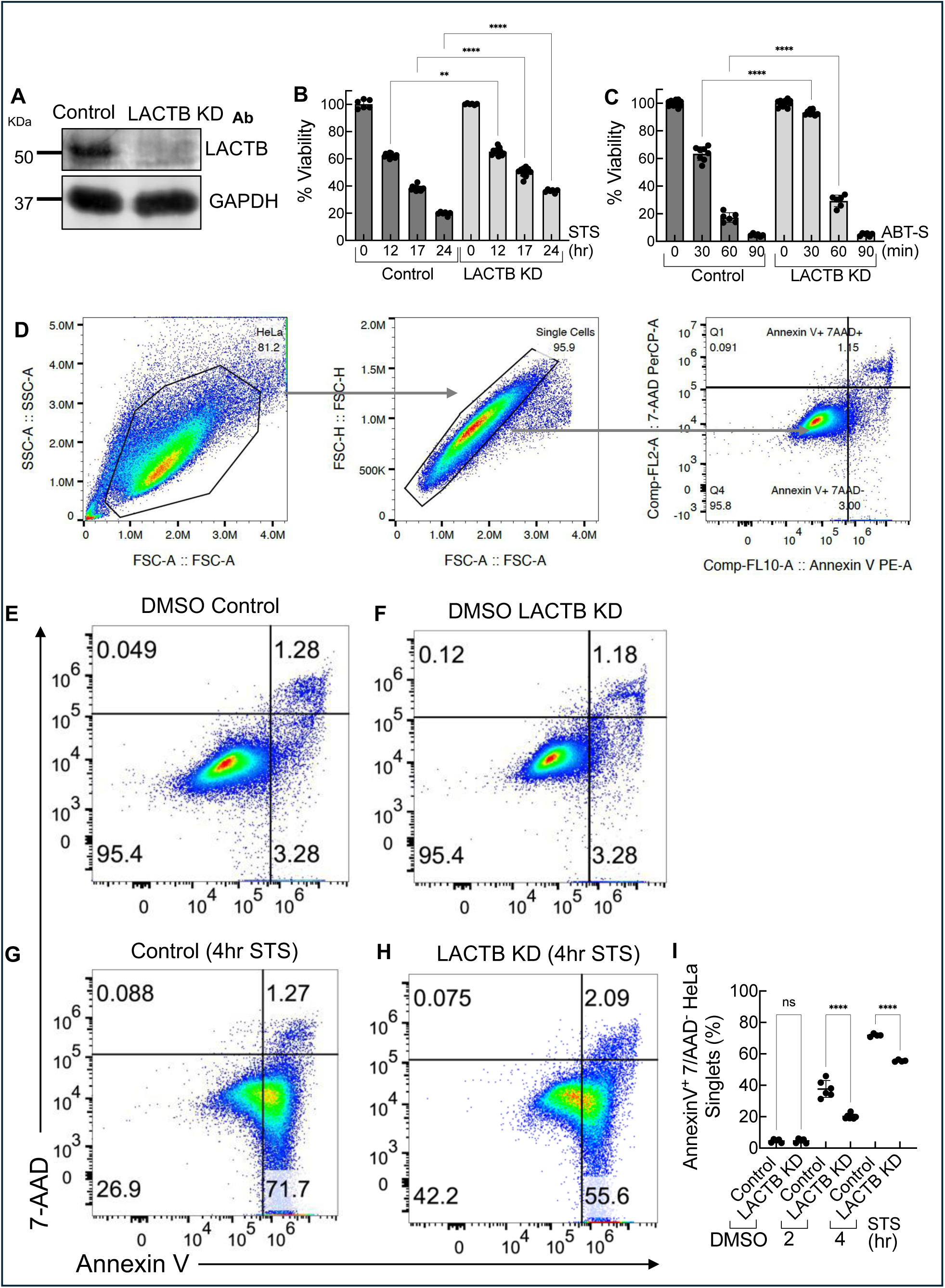
LACTB knockdown reduces apoptosis. (**A**) Western blot for LACTB in control or LACTB knockdown (KD) B16-F10 cells. GAPDH used as loading control. (**B**) Quantification of SRB assay in control and LACTB KD B16-F10 cells treated with 1 μM staurosporine (STS) for various times. N = 2. (**C**) Quantification of SRB assay in control and LACTB KD B16-F10 cells treated with ABT-S for various times. N = 2. (**D**) Representative gating strategy used for flow cytometry analysis of HeLa cells for AnnexinV and 7-AAD staining. Cells were initially gated based on forward scatter area (FSC-A) versus side scatter area (SSC-A) to identify the population of interest, followed by FSC-A versus forward scatter height (FSC-H) to exclude doublets. (**E**, **F**) Annexin-V and 7-AAD staining of control (E) and LACTB KD (F) HeLa cells treated with DMSO for 4 hours.. (**G**, **H**) Annexin-V and 7-AAD staining of control (G) and LACTB KD (H) HeLa cells treated with 1 μM STS for 4 hours. **(I)** Quantification of Annexin-V^+^/7^-^AAD -HeLa singlets from three independent control and LACTB KD experiments upon STS treatment. ****p<0.0001, **p<0.01 (one-way ANOVA). Data are presented as mean ± SD. N represents the number of independent experiments.

**Figure S2:**
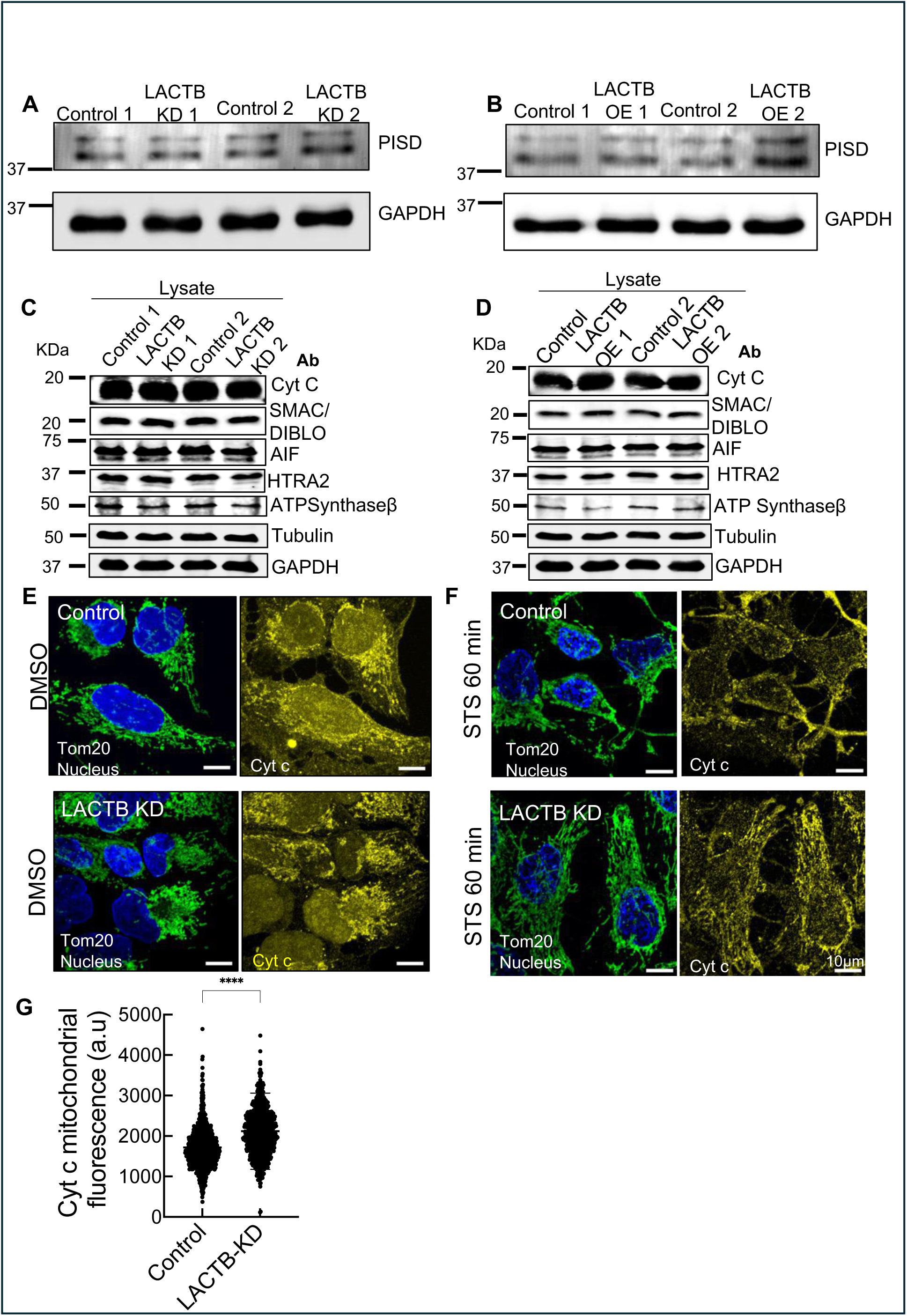
LACTB knockdown and overexpression do not alter expression levels of mitochondrially-released factors or PISD. (**A,B**) PISD expression levels in control and LACTB KD (A; two independent knockdowns) or control and LACTB OE HeLa cells (B; two independent overexpression lines). (**C,D**) Western blot analysis of total cell extracts from control and LACTB KD (C) or control and LACTB OE (D) HeLa cells for cytochrome c and other mitochondrial proteins. GAPDH and tubulin used as loading controls. Experiments were performed using two independent LACTB KD samples or LACTB OE lines. (**E,F**) Immunofluorescence staining for cytochrome c (yellow) and Tom20 (green) in U2-OS cells under DMSO treatment (E) or 1 μM STS for 1 hour (F). Blue, DAPI. (**G**) Quantification of mitochondrial cytochrome c levels upon 1-hour STS treatment in U2-OS cells (N_mito_ ≥ 735, N_cells_ = 15) ****p < 0.0001 (Mann–Whitney test). Data are presented as mean ± SD.

**Figure S3:**
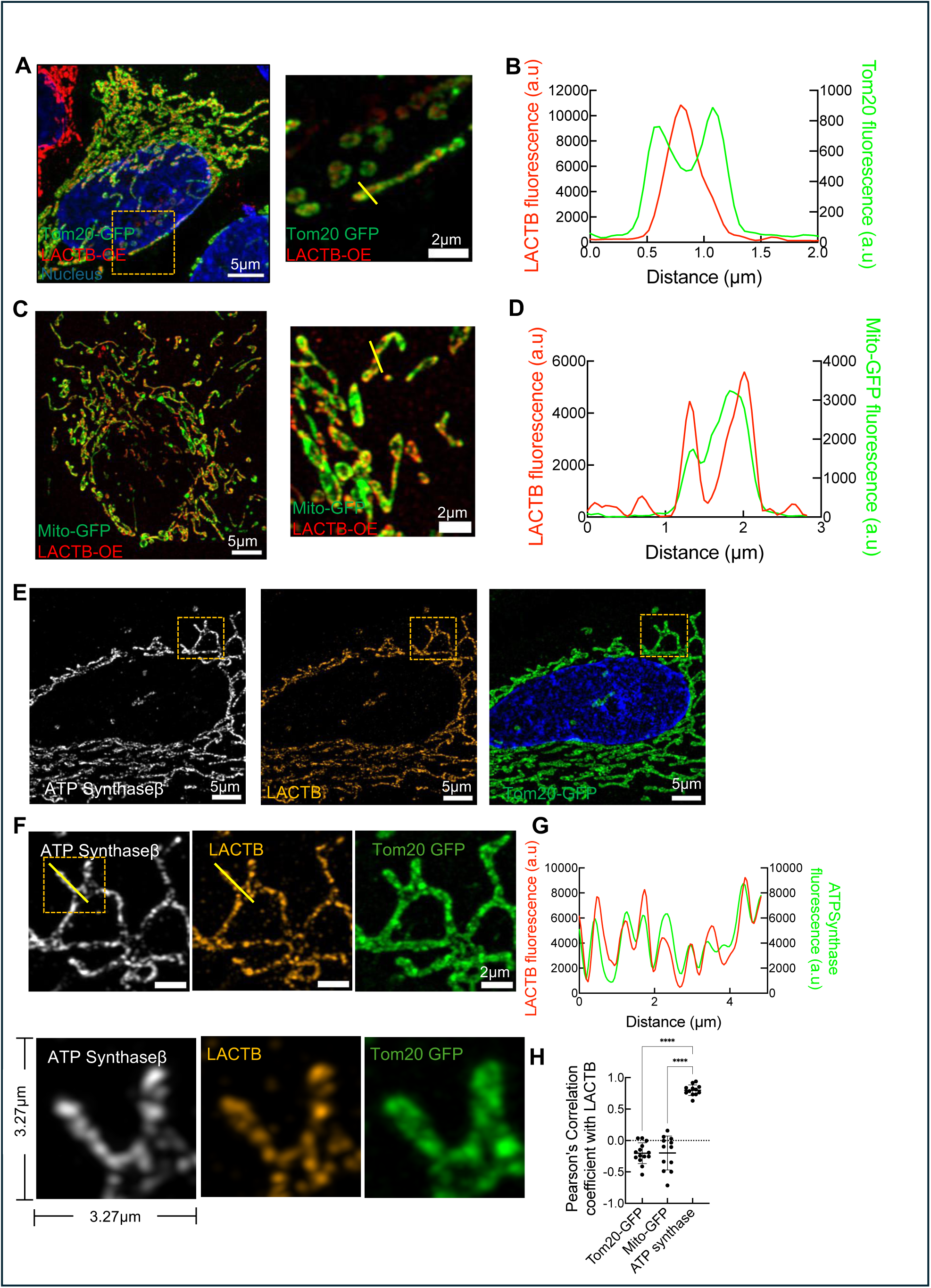
LACTB colocalizes with ATP Synthase in mitochondria. (**A**) Immunofluorescence staining of LACTB (red) in LACTB-OE HeLa cells transiently transfected with Tom20-GFP (green). DAPI, blue. Image on the right shows zoom of the boxed region. (**B**) Line profile through a mitochondrion in panel (A). (C) Immunofluorescence staining of LACTB (red) in LACTB-OE HeLa cells transiently transfected with Mito-GFP (green). DAPI, blue. Image on the right shows zoom of the boxed region. (**D**) Line profile through a mitochondrion in panel (C). (**E**) Immunofluorescence staining LACTB (orange) and ATP synthase b (gray) in LACTB-OE HeLa cells transiently expressing Tom20-GFP (green). DAPI, blue. (**F**) Zoomed image from box in panel (E), comparing localization pattern of ATP synthase b, LACTB and Tom20. Images below show a further zoom of the mitochondrion in the line profile analysis in panel G. (**G**) Line profile analysis of LACTB and ATP synthase b signals from panel (F). (**H**) Pearson’s colocalization coefficient for colocalization between LACTB and Tom20-GFP, Mito-GFP, or ATP Synthase. ****p < 0.0001 (one-way ANOVA). Data are presented as mean ± SD. N_mito_ ≥ 12.

**Figure S4:**
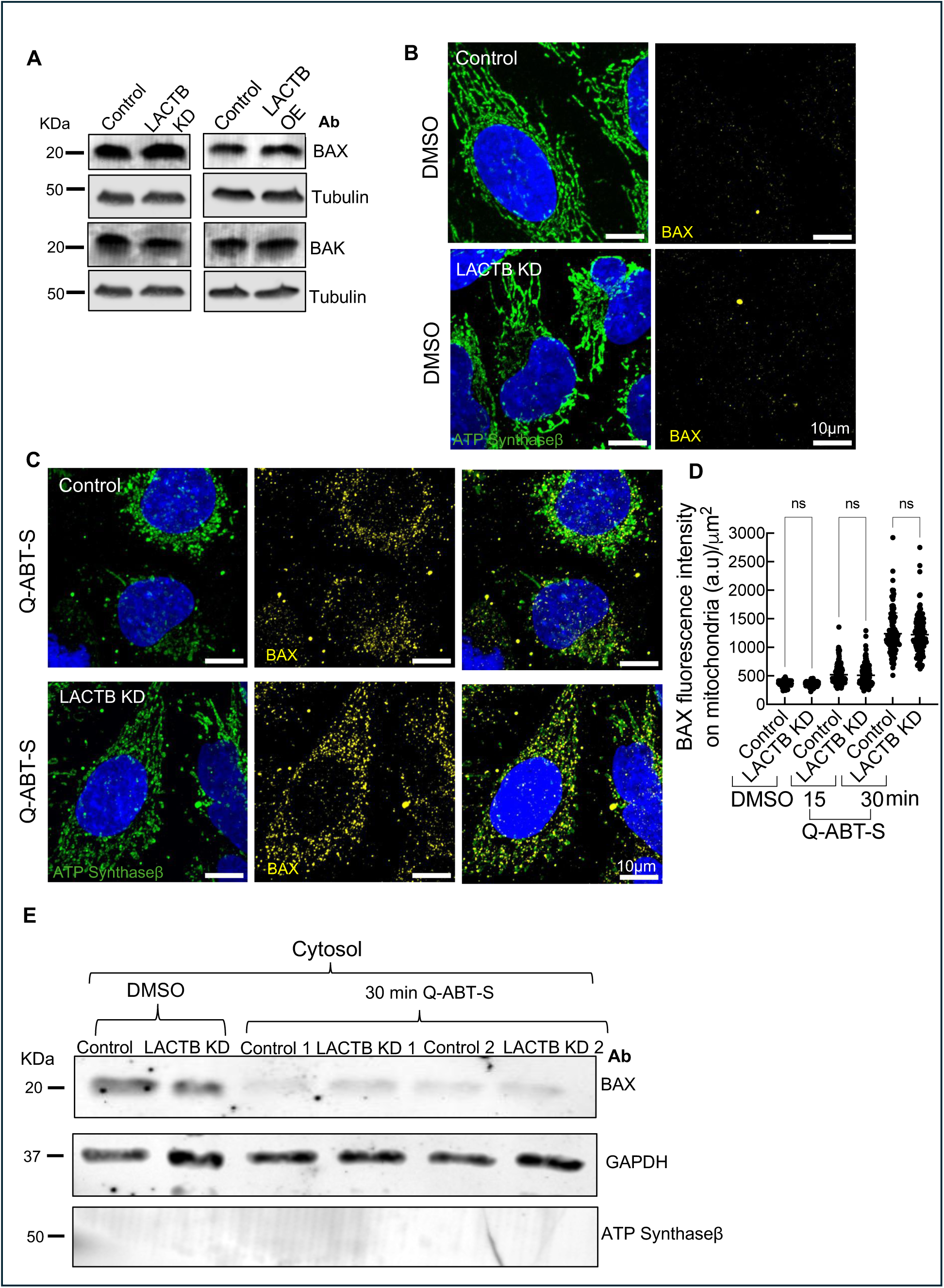
LACTB does not influence BAX recruitment to mitochondria. (**A**) Western blot analysis of BAX and BAK levels in total cell extracts from control, LACTB KD or LACTB OE HeLa cells. Tubulin used as a loading control. (**B**, **C**) Immunofluorescence staining of endogenous BAX (yellow) in control and LACTB KD HeLa cells treated with DMSO (B) or ABT-S (C) for 30 min. Prior to ABT-S treatment, the pan-caspase inhibitor Q-VD-OPh (20 μM) was added for 1-hour. ATP Synthase staining for mitochondria in green, and DAPI staining for nuclei in blue. BAX signal intensity was brightness and contrast adjusted to specifically highlight its mitochondrial localization. (**D**) Quantification of mitochondrial BAX fluorescence intensity (intensity per μm² mitochondrial area) in control and LACTB KD HeLa cells treated with Q-ABT-S for 15 or 30 minutes. Data represent mean ± SD from 2 independent experiments, with N_cells_ ≥ 104. Statistical analysis using one-way ANOVA indicates no significant difference (ns). (**E**) Western blot analysis of BAX, GAPDH, and ATP synthase b in cytosolic extracts from control and LACTB KD HeLa cells after 30 min DMSO or Q-ABT-S treatment. Experiment conducted with two independent LACTB KD samples.

**Figure S5:**
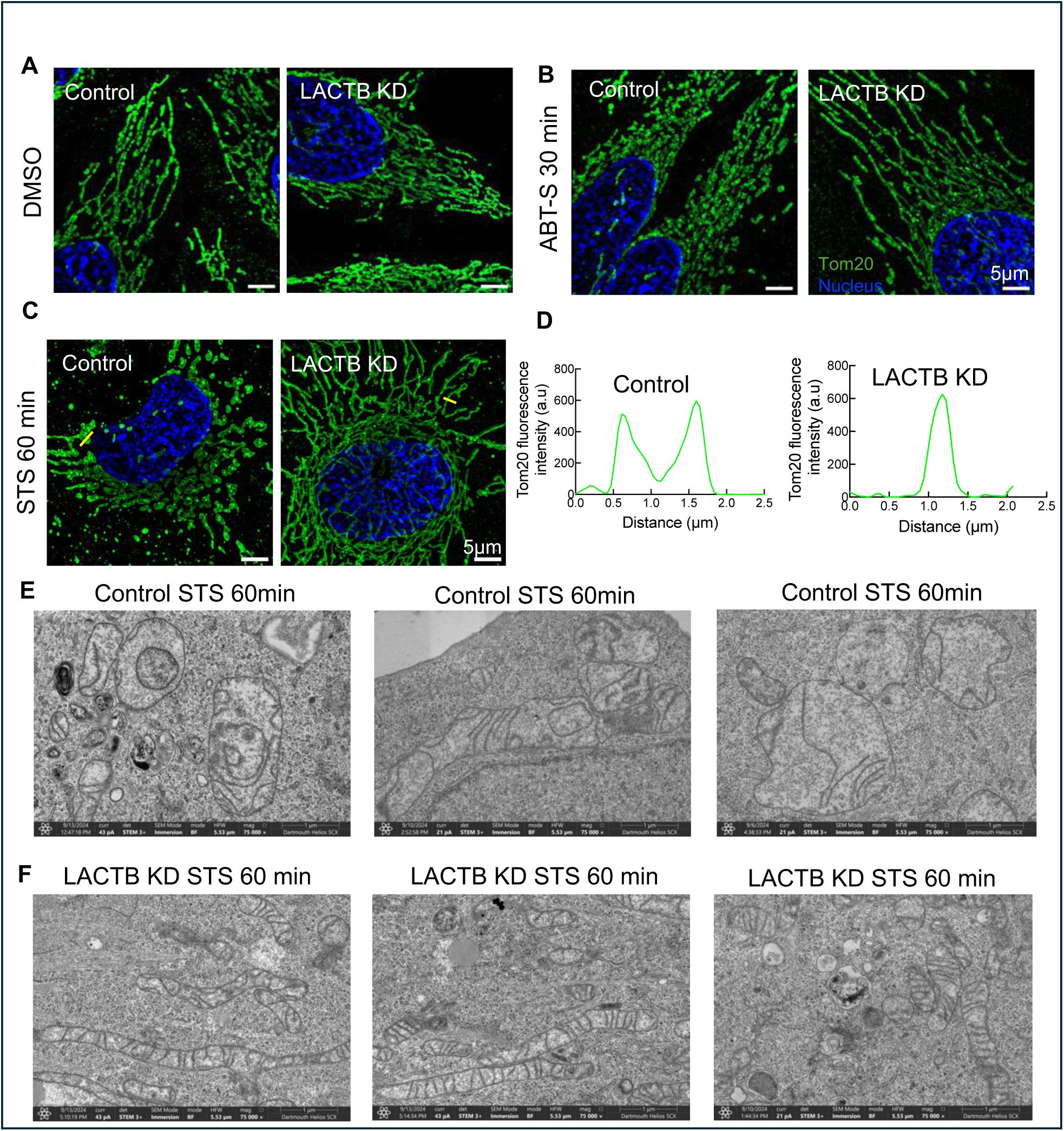
LACTB mediates mitochondrial remodelling upon apoptosis induction. (**A**-**C**) Full images of magnified insets from Fig 3A-C showing immunofluorescence staining of Tom20 (green) in control and LACTB KD HeLa cells treated with 1 hr DMSO (A), 30 min ABT-S (B) or 1 hr STS (C). (**D**) Line profile analysis of Tom20 fluorescence intensity in control and LACTB KD HeLa cells treated with STS for 1 hour. Lines shown in panel C. (**E**, **F**) TEM micrographs of mitochondria in control (E) and LACTB KD (F) HeLa cells treated with STS for 60 min.

**Figure S6.**
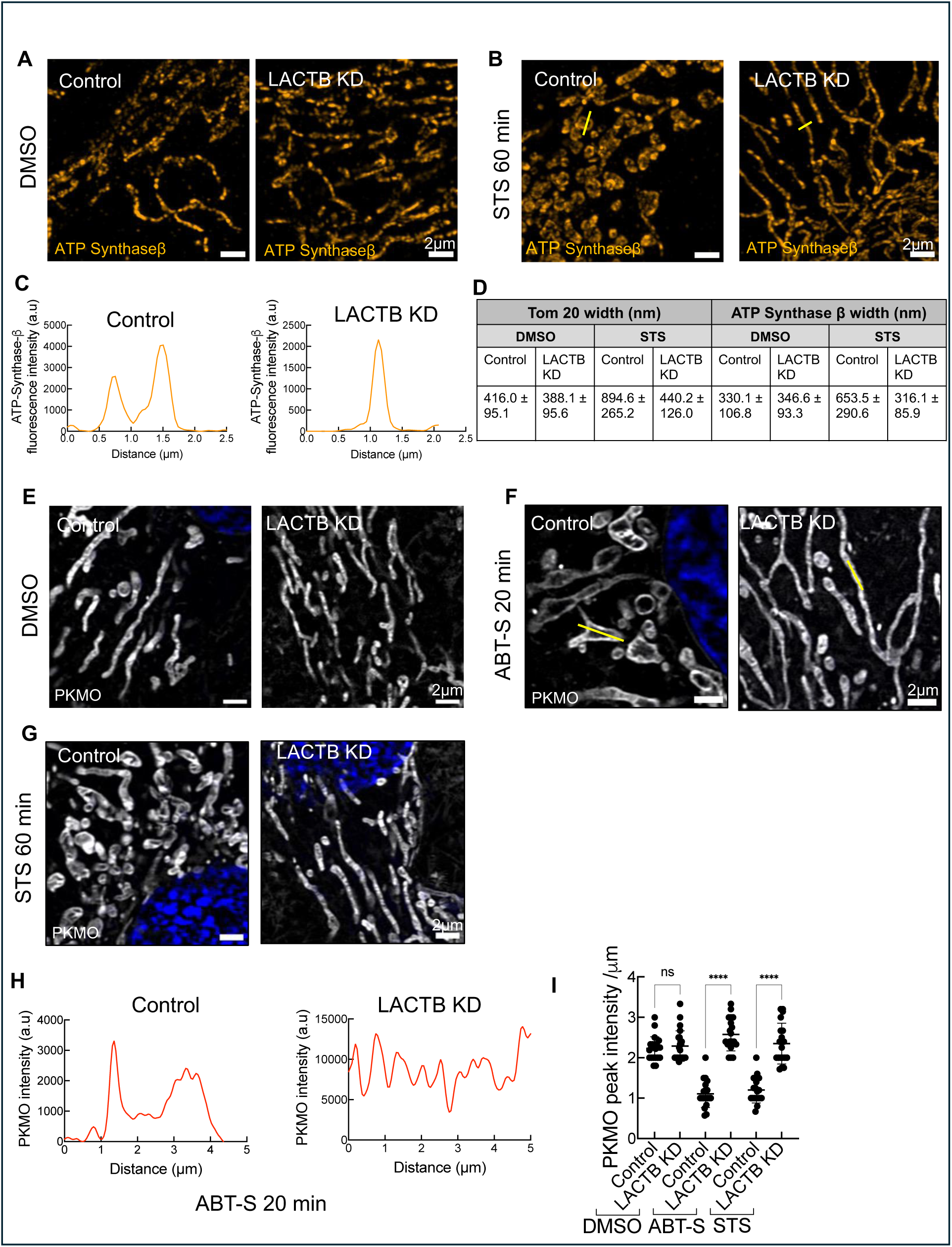
LACTB KD preserves IMM morphology during apoptosis. (**A**,**B**) Immunofluorescence staining of ATP Synthase β (orange) in control or LACTB KD HeLa cells upon DMSO (A) or staurosporine (STS) treatment (B) for 60 min. (**C**) Line scan analysis of ATP Synthase β fluorescence intensity in control and LACTB KD HeLa cells treated with STS for 60 min. (**D**) Table summarizing mitochondrial widths (mean ± SD, in nm), as measured by Tom20 or ATP synthase b markers, of control or LACTB KD HeLa cells upon DMSO or STS treatment. (**E**-**G**) PKmito Orange FX (PKMO) staining (cristae marker, in grey) in control siRNA and LACTB KD HeLa cells upon DMSO (E), 20 min ABT-S (F), or 1-hr STS (G) treatment. Blue, DAPI. (**H**) Line scan analysis of PKMO fluorescence intensity in control and LACTB KD cells treated with ABT-S for 20 min. (Characterization of membrane binding) Quantification of PKMO intensity peaks (per micron) using line profile analysis of mitochondria. N_mito_ ≥ 19. Error bars represent mean ± SD. ****p < 0.0001; ns, not significant (one-way ANOVA).

**Figure S7:**
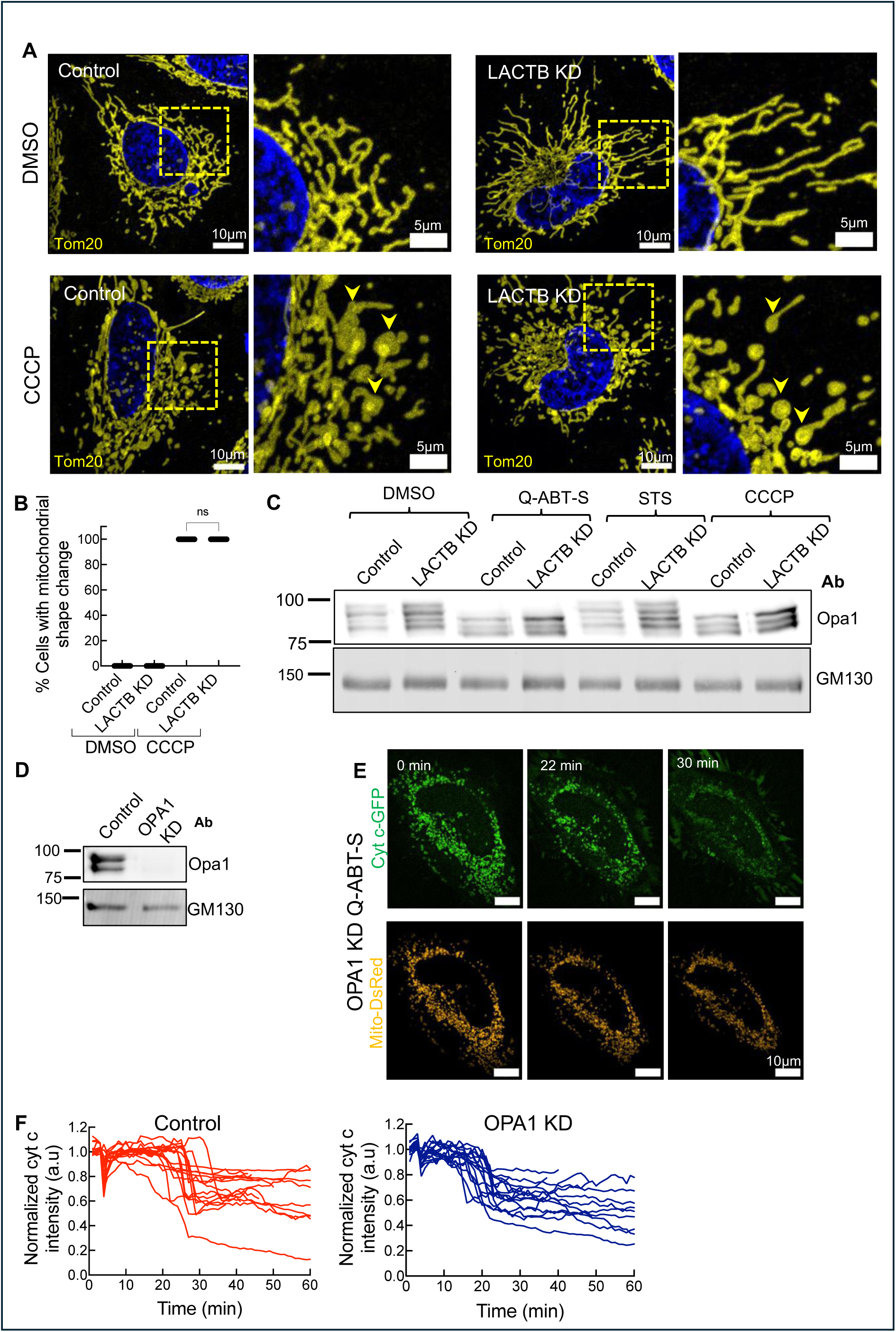
LACTB action is independent of OPA1. (**A**) Immunofluorescence staining of Tom20 (yellow) in control siRNA and LACTB KD HeLa cells after 20 min DMSO or CCCP (20 μM) treatment. Blue, DAPI. Panels to right are zooms of boxed regions. (**B**) Scatter plot showing blinded analysis of cells undergoing mitochondrial shape change (assessed by Tom20 immunofluorescence) in control siRNA and LACTB KD HeLa cells after 20 min of DMSO or CCCP (20 μM) treatment. N_cells_ ≥ 24. Error bars represent mean ± SD. ****p < 0.0001; ns, not significant (one-way ANOVA). (**C**) Western blot assessing OPA1 processing in control and LACTB KD HeLa cells under DMSO (1hour), Q-ABT-S (30min), STS (1hour), and CCCP (20 min) treatments. GM130, loading control. (**D**) Western blot of OPA1 in control and OPA1 KD HeLa cells. GM130, loading control. (**E**) Live-cell imaging of GFP-tagged cytochrome c release in OPA1 KD HeLa cells treated with Q-ABT-S for the indicated times. (**F**) Quantification of cytochrome c release kinetics after Q-ABT-S treatment in control and OPA1 KD HeLa cells. N = 3 independent experiments; N_cells_= 16.

**Figure S8:**
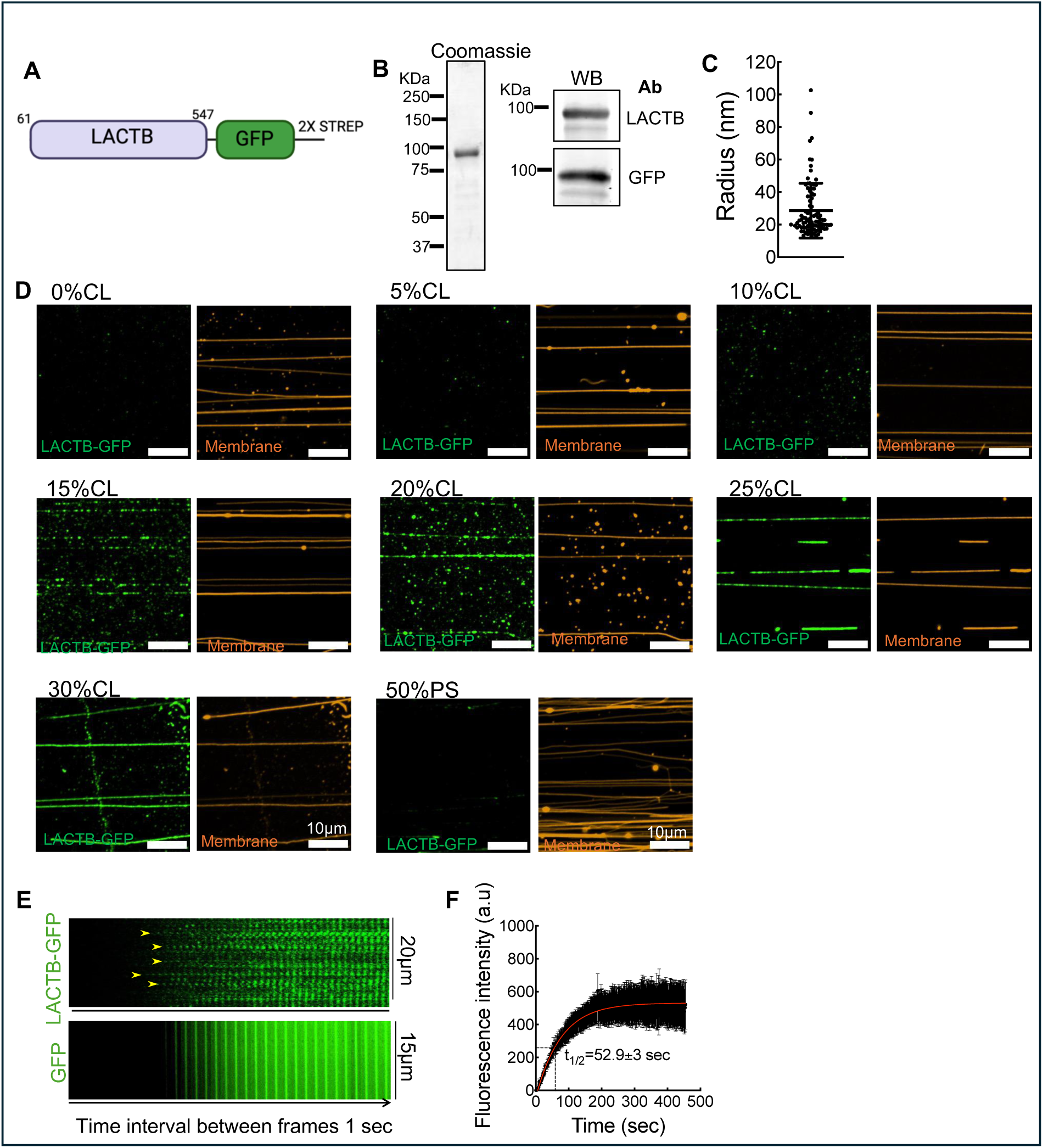
Characterization of membrane binding by LACTB. (**A**) Cartoon representation of the LACTB construct used for biochemical studies. The N-terminal 60 amino acids are LACTB’s mitochondrial targeting sequence, and were removed from the construct. 2x STREP = tandem Strep purification tag. (**B**) Coomassie-stained SDS-PAGE gel and western blot of purified LACTB-GFP. (**C**) Distribution of radii for membrane nanotubes in the system used here, computed using a previously described method(*43*, *44*). Error bars represent mean ± SD. (**D**) Representative micrographs of LACTB-GFP binding to membrane nanotubes (10 min incubation) as a function of cardiolipin concentration. Lipid mixtures contain PC and 0.5 mol% RhPE. Varying % of CL or phosphatidylserine (PS) are added (with corresponding % of PC removed). (**E**) Representative timelapse montage showing nucleation (yellow arrowheads) and growth of LACTB-GFP scaffolds on nanotubes. Control experiments with uniform recruitment of 6His-GFP (onto nanotubes containing Ni-NTA lipid) were also performed. Lipid compositions: DOPC:CL:RhPE (74.5:25:0.5 mol%) for LACTB-GFP; DOPC:PS:DGS-NiNTA:RhPE (79.5:15:5:0.5 mol%) for 6His-GFP. (**F**) Rate of LACTB-GFP binding to nanotubes. N = 2 experiments; n_tubes_ =10

**Figure S9:**
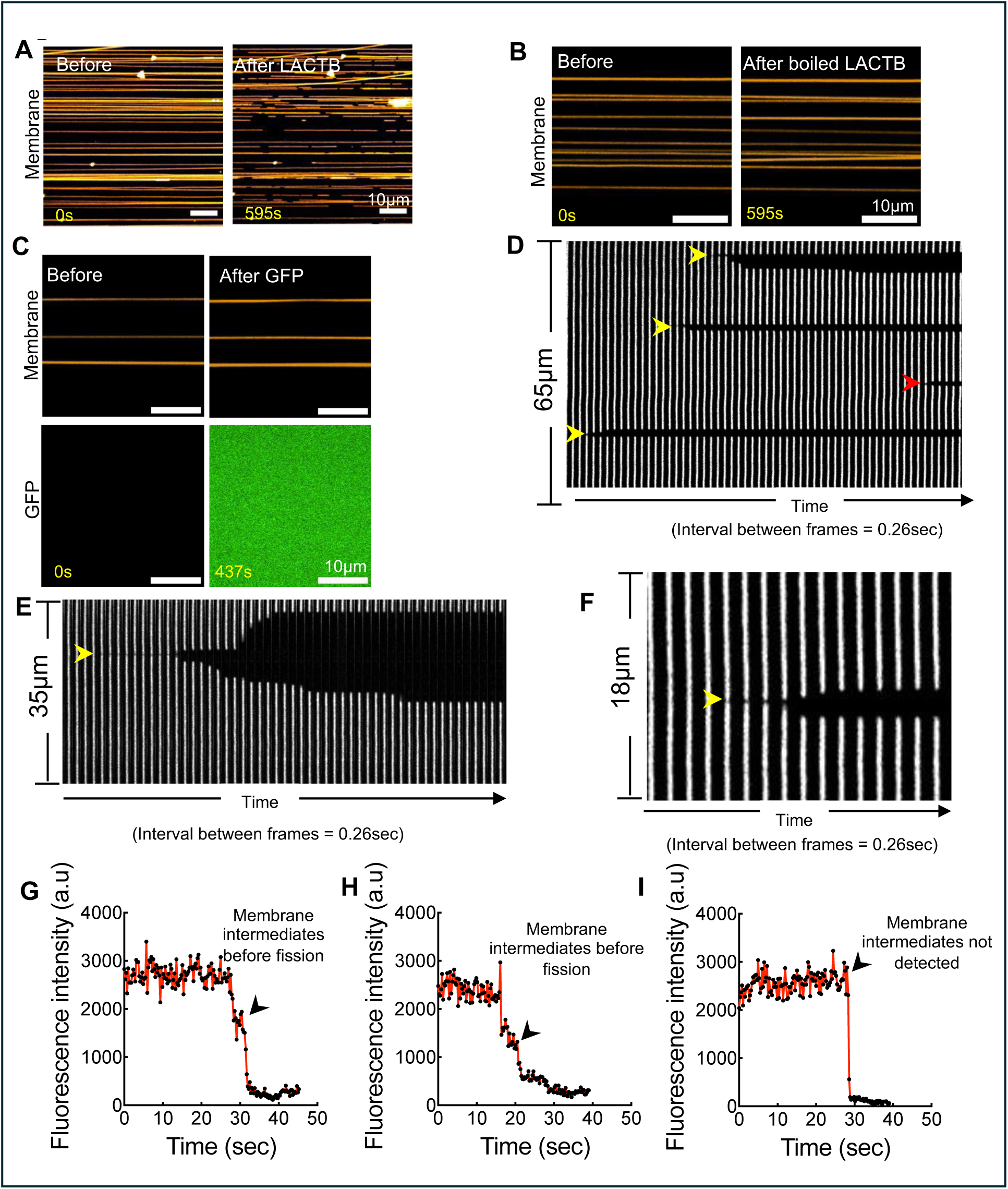
LACTB-mediated membrane fission. (**A**) Full imaging field from which the inset in Figure 5i was taken, before and 10 min after flowing LACTB-GFP onto nanotubes containing 25 mol% cardiolipin. (**B**) Representative images showing membrane nanotubes containing 25 mol% cardiolipin before and after flowing in boiled LACTB-GFP (10min incubation). (**C**) Representative images showing membrane nanotubes containing 25 mol% cardiolipin before and after flowing 1 μM GFP. (**D**–**F**) Representative kymographs of nanotubes during LACTB-mediated remodelling, showing fission events (arrowheads). Constriction is marked by a decrease in fluorescence intensity prior to fission (indicated by yellow arrowheads). A fission event without detectable prior constriction is also shown (red arrowhead). (**G**–**I**) Single-pixel fluorescence traces extracted from kymographs highlighting the kinetics of membrane nanotube constriction before fission events. Panel (G) and (H) show two examples of constriction before fission, while panel (I) shows a fission event without detectable prior constriction. Fission event marked by black arrow head.

**Table S1.** Details of antibodies used in this study, including sources, catalog numbers, and dilutions.

| **Antibody** | **Source** | **Catalog number** | **Dilution** |
| --- | --- | --- | --- |
| Rabbit anti-LACTB | Proteintech | 18195-1-AP | 1:1000 WB; 1:100 IF |
| Rabbit anti-PISD | Proteintech | 16401-1-AP | 1:500 WB |
| Mouse anti-Cytochrome c | Abcam | 110325 | 1:1000 WB; 1:200 IF |
| Rabbit anti-BAX | Cell Signaling Technology | 2772T | 1:1000 WB; 1:200 IF |
| Rabbit anti-BAK | Cell Signaling Technology | 12105T | 1:1000 WB |
| Rabbit anti-HTRA2/Omi | Proteintech | 15775-1-AP | 1:1000 WB |
| Rabbit anti-SMAC/DIABLO | Proteintech | 10434-1-AP | 1:500 WB |
| Rabbit anti-AIF | Proteintech | 67791-1-Ig | 1:1000 WB |
| Mouse anti-ATP Synthase $\beta$ | Invitrogen (Molecular Probes) | A21351 | 1:1000 WB; 1:200 IF |
| Rabbit anti-Tom20 | Abcam | ab78547 | 1:1000 WB; 1:200 IF |
| Mouse anti-GAPDH | Santa Cruz | sc-365062 | 1:2000 WB |
| Rabbit anti-GM130 | Transduction labs | G65120 | 1:1000 WB |
| Mouse anti-OPA1 | BD Biosciences | 612606 | 1:2000 WB |
| Mouse anti-Tubulin | Sigma-Aldrich | T9026, mouse, clone DM1-α | 1:4000 WB |
| Goat anti-rabbit IgG Texas Red secondary | Vector Laboratories | TI-1000 | 1:500 IF |
| Horse anti-mouse IgG fluorescein secondary | Vector Laboratories | FI-2000 | 1:500 IF |
| Goat anti-rabbit IRDye 800CW | LICOR | 926-32211 | 1:15,000 WB |
| Goat anti-mouse<br>IRDye 680RD | LICOR | 926-68070 | 1:15,000 WB |
| Goat anti-mouse IgG<br>HRP conjugate | Bio-Rad | 1705047 | 1:2000 WB |
| Goat anti-rabbit IgG<br>HRP conjugate | Bio-Rad | 1706515 | 1:5000 WB |

### Supplementary Movies

**Supplementary Movie 1**

Time-lapse imaging of cytochrome c release in control cell upon apoptosis induction. Control siRNA-treated cell expressing cytochrome c-GFP (green) was treated at 4 min with ABT-S (10 μM ABT-737 and 2 μM S63845) in the presence of the pan-caspase inhibitor Q-VD-OPh (20 μM, 1-hour preincubation). Scale bar: 10 μm. Time interval: 1 min.

**Supplementary Movie 2**

Mitochondrial shape change in control cell upon apoptosis induction. Control siRNA-treated cell expressing Mito-dsRed (orange) was treated as described above. Scale bar: 10 μm. Time interval: 1 min.

**Supplementary Movie 3**

Cytochrome c kinetics in LACTB knockdown (KD) cell upon apoptosis induction. LACTB KD cell expressing cytochrome c-GFP (green) was treated at 4 min with Q-ABT-S as described above. Scale bar: 10 μm. Time interval: 1 min.

**Supplementary Movie 4**

Mitochondrial shape change in LACTB KD cell upon apoptosis induction. LACTB KD cell expressing cytochrome c-GFP (green) was treated as described above. Scale bar: 10 μm. Time interval: 1 min.

**Supplementary Movie 5**

Time-lapse imaging of LACTB (1 μM) catalysing fission of membrane nanotubes containing 25 mol% cardiolipin. Scale bar: 10 μm. Time interval: 0.27 sec.

